# Alternative NADH:ubiquinone oxidoreductase modulates disease susceptibility in rice by interfering ROS homeostasis and ferroptosis

**DOI:** 10.64898/2025.12.08.692912

**Authors:** Debashis Sahoo, Ravindra K Chandan, Naveen Goel, Gopaljee Jha

**Author notes:** to whom correspondence should be addressed, **Corresponding author information:** Gopaljee Jha, PhD.

## Abstract

Necrotrophic fungal pathogens, such as *Rhizoctonia solani*, the causal agent of sheath blight disease (SBD) in rice, enhance the production of reactive oxygen species (ROS) to induce necrosis and infect a broad range of plant species. Here, we present evidence that the upregulation of host alternative NADH:ubiquinone oxidoreductase (OsNUOR) is important for *R. solani* to induce ROS production and cause SBD in rice. The knock-out lines developed through genome editing are defective in ROS accumulation and demonstrate SBD resistance, whereas the overexpression lines exhibit enhanced disease susceptibility. We emphasize that the cross-talk between OsNUOR and ROS signaling is important for the suppression of antioxidant defense, followed by enhancement of lipid peroxidation and accumulation of ferric ions in rice under *R. solani*-infected conditions. Considering that treatment with deferoxamine, an iron chelator, and ferrostatin-1, a lipid peroxidation inhibitor, prevents SBD disease, whereas FeCl_3_ enhances disease severity, we propose that *R. solani* induces iron-dependent cell death, referred to as ferroptosis, in rice. As agronomic traits of OsNUOR-edited lines are comparable to wild-type lines, we emphasize that editing of OsNUOR and additional genes in ferroptotic pathway can be utilized as biotechnological interventions for control of SBD, for which effective control measures remain a challenge.

## Introduction

Sheath blight disease (SBD), caused by a necrotrophic fungal pathogen *Rhizoctonia solani* AG1-IA, is one of the deadly fungal diseases of rice that accounts for 20-42% annual yield loss ^1^. Due to a lack of resistance sources, the disease is mostly managed by excessive use of fungicides, which not only add costs to rice cultivation but also poses a concern for consumers as well as the environment ^2,3^. Therefore, efforts are being made to understand the molecular intricacies of SBD in rice and identify key players that modulate disease susceptibility. Pathological studies have emphasized that *R. solani* undergoes a necrotrophic phase wherein it causes the disintegration of host cells, culminating in visual necrotic disease lesions ^4^. The transcriptome, metabolome, and proteome-based studies have revealed that photosynthetic machinery is downregulated, while respiratory as well as secondary metabolic pathways are enhanced during *R. solani*-induced necrotrophy in rice ^4^. In addition, the downregulation of SA-based defense ^5,6^ and enhanced accumulation of reactive oxygen species (ROS) are positively associated with disease severity ^4,7^. The host functions that are modulated during *R. solani* infection to facilitate an oxidative stress-enriched environment and enhance disease susceptibility, remain to be elucidated.

In eukaryotic cells, mitochondria are major sites for ROS production, wherein complex-I (NADH:ubiquinone oxidoreductase, NUOR) and III (cytochrome bc1 complex; ubiquinol:cytochrome c oxidoreductase) of electron transport chain (ETC) contribute to the production of superoxides and other forms of ROS ^8,9^. The complex-I has rotenone-sensitive NUOR activity, wherein NADH dehydrogenase module (N-module) catalyzes NADH oxidization and passes electrons into its Fe-S cluster containing subunits. The electrons are subsequently routed through other complexes of ETC for ATP generation ^10^. Plants and fungi encode mitochondrial localized rotenone-insensitive alternative NADH:ubiquinone oxidoreductase (alternative NUOR) proteins that due to NADH dehydrogenase activity can feed electrons (similar to complex-I) into ETC ^11^. The complex-I has a proton-pumping P-module that creates a transmembrane proton gradient to drive ATP generation. As alternate NUORs lack a proton-pumping P-module, they do not contribute to ATP generation ^12,13^. Our previous study reported that alternative *NUOR* (rice: LOC_Os07g37730 and tomato: XM_004232770) is a potential host susceptibility factor in rice/tomato ^14^. It gets upregulated upon *R. solani* infection, and the knock-down of its expression imparts disease tolerance. In the present study, we demonstrate that overexpression of *OsNUOR* enhances disease susceptibility, whereas loss of function alleles created by CRISPR-Cas9-based genome editing imparts disease tolerance in rice. We present evidence that OsNUOR is important for ROS accumulation, lipid peroxidation, and iron accumulation in rice under infected conditions. Moreover, we emphasize that necrotic symptoms associated with SBD are due to iron-dependent cell death referred to as ferroptosis ^15,16^. The role of OsNUOR in promoting disease susceptibility through modulation of redox-signaling and ferroptosis is described.

## Results

### *OsNUOR* edited lines exhibit sheath blight disease resistance, without compromising agronomic traits

To investigate the role of *OsNUOR* during *R. solani* infection, we generated overexpression (OE) lines under CaMV35S promoter (Supplementary Figures 1A-D) and SDN-1 type genome-edited lines using CRISPR-Cas9 approach (Supplementary Figure 2A-D) in rice (cv. TP309). Based on the extent of *OsNUOR* upregulation (Supplementary Figure 1E), two independent OE (OE3 and OE5) lines were selected for further analysis at T_2_ generation. We selected two of the edited lines (m6 and m10) having downregulated expression of *OsNUOR* (Supplementary Figure 2E) for further analysis at T_2_ generation (Figures 1A and 1B). The m6 line has 272-325 bp deletion (91-108 aa), while m10 line has 271-321 bp (91-107 aa) deletion in the exonic region. In addition, both the lines have nucleotide substitution at 262-264 bp, which resulted in the substitution of proline (P) (88^th^ aa) to asparagine (N). The functional domain of OsNUOR, i.e pyridine nucleotide-disulfide oxidoreductase (Pyr_redox2 domain), belonging to NADH dehydrogenase family remained intact in edited lines (Supplementary Figure 3). BLAST searches identified an additional paralog of *OsNUOR* (LOC_Os01g61410) (Supplementary Figure 4A) having Pyr_redox2 domain (Supplementary Figure 4B). However, the gene was only marginally expressed under *R. solani*-infected conditions in different rice lines (Supplementary Figure 4C).

**Figure 1:**
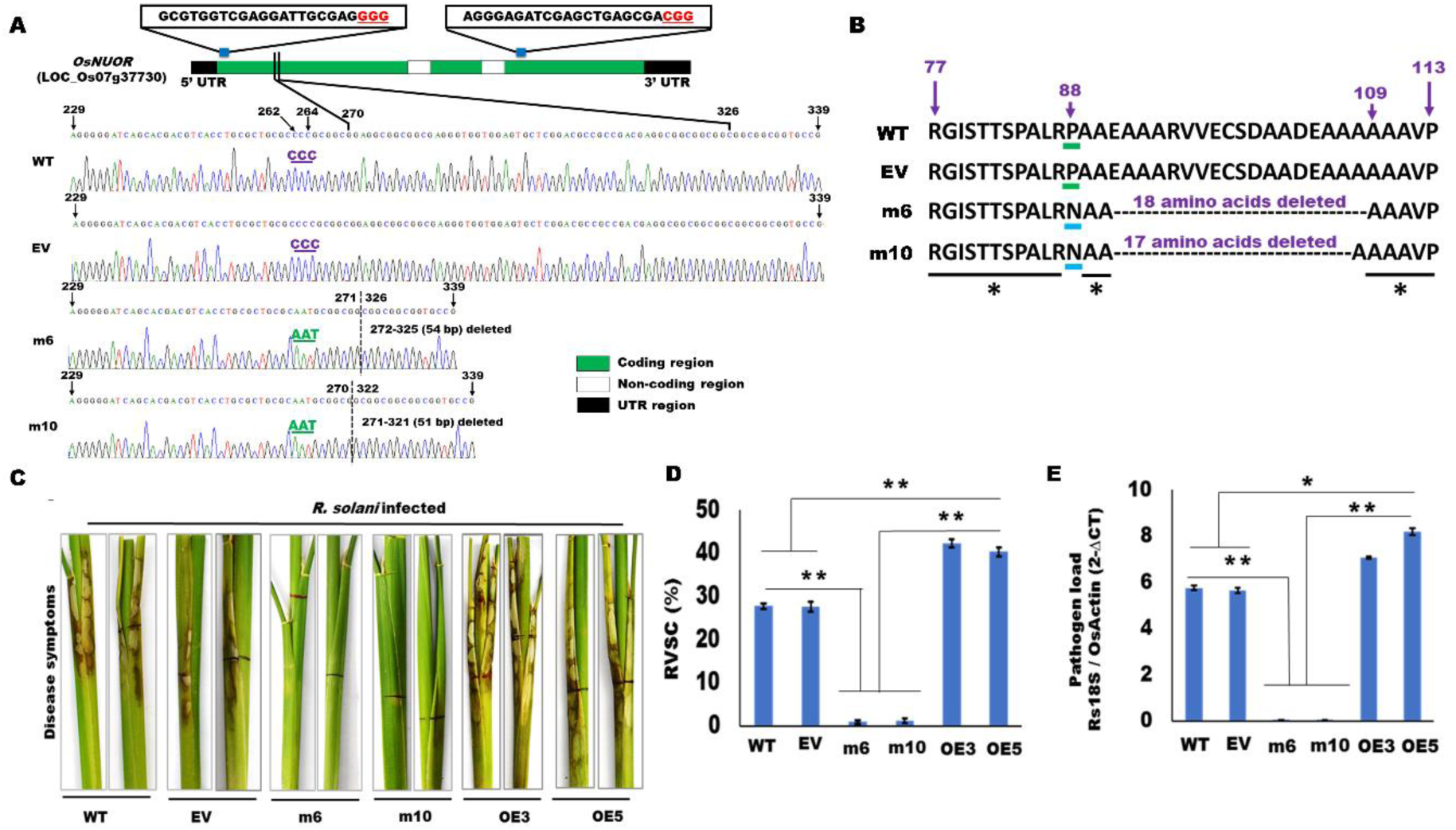
CRISPR-Cas9 mediated genome editing of *OsNUOR* imparts sheath blight disease tolerance in rice. **(A)** The schematics represent the position of the guide RNAs (sequence provided in the box), the corresponding PAM sequences (in red), and the target region of OsNUOR in rice lines. The chromatograms of the edited region have been depicted. The dotted line represents the location of the 54 bp and 51 bp deletion in the edited lines, m6 and m10, respectively. At 262-264 bp region, there is a substitution of CCC in WT (marked in violet) to AAT in the edited lines (marked in green). **(B)** Amino acid sequence alignment, showing 91 to 108 aa deletion in m6, and 91 to 107 aa deletion in m10 lines. Proline at 88 aa positions in WT is converted to asparagine in the edited lines. The asterisk (*) represents consensus amino acid residues of OsNUOR in different rice lines. **(C)** Disease symptoms in WT, EV, edited, and overexpression (OE3 and OE5) lines upon *R. solani* infection, at 5 dpi. Bar graph showing **(D)** Relative vertical sheath colonization (RVSC) based disease index and **(E)** qRT-PCR based quantification of *R. solani* biomass in different rice lines. The abundance of *18S rRNA* of *R. solani* is measured using ΔCt method, where ΔCt is the difference between the Ct value of *18S rRNA* and *Oryza sativa actin* gene. The average of three independent biological replicates is plotted and the error bar represents mean ± standard error. ‘*’ and ‘**’ indicate significant differences at *p* < 0.05 and *p* < 0.001, respectively (estimated using one-way ANOVA and Student-Newman-Keuls test).

It was noteworthy that compared to WT (wild-type) and EV (empty vector control) lines, *OsNUOR*-edited lines (m6 and m10) had significantly reduced disease symptoms, whereas OE lines (OE3 and OE5) had enhanced disease symptoms (Figure 1C). Relative vertical sheath colonization (RVSC) based disease index (Figure 1D) and pathogen load (estimated as an abundance of 18S rRNA of *R. solani* by qRT-PCR, reflecting the abundance of fungal biomass) (Figure 1E) were also significantly reduced in the edited lines, but enhanced in OE lines, compared to WT and EV plants. The agronomic traits of *OsNUOR-*edited and OE lines were comparable to WT/EV lines, under controlled conditions (Supplementary Figures 5A-D).

### OsNUOR localizes in mitochondria and interacts with 75 kDa subunit of complex 1

The targetP (https://services.healthtech.dtu.dk/services/TargetP-2.0/) analysis suggested that OsNUOR is localized in mitochondria (Supplementary Table 1). This was reinforced by reporter assay, which revealed that the yellow signal of YFP-tagged OsNUOR coincides with the red signal of the mitochondria-mCherry marker, CD3-991 ^17^ in *Nicotiana benthamiana* leaves (Figure 2A). Considering that both the edited lines had nucleotide alterations at the N-terminal region, we deleted 1-108 aa residues, created a variant of OsNUOR (OsNUOR_1), and analyzed its subcellular localization. The reporter assay revealed that OsNUOR_1 is not localized in mitochondria (Figure 2A).

**Figure 2:**
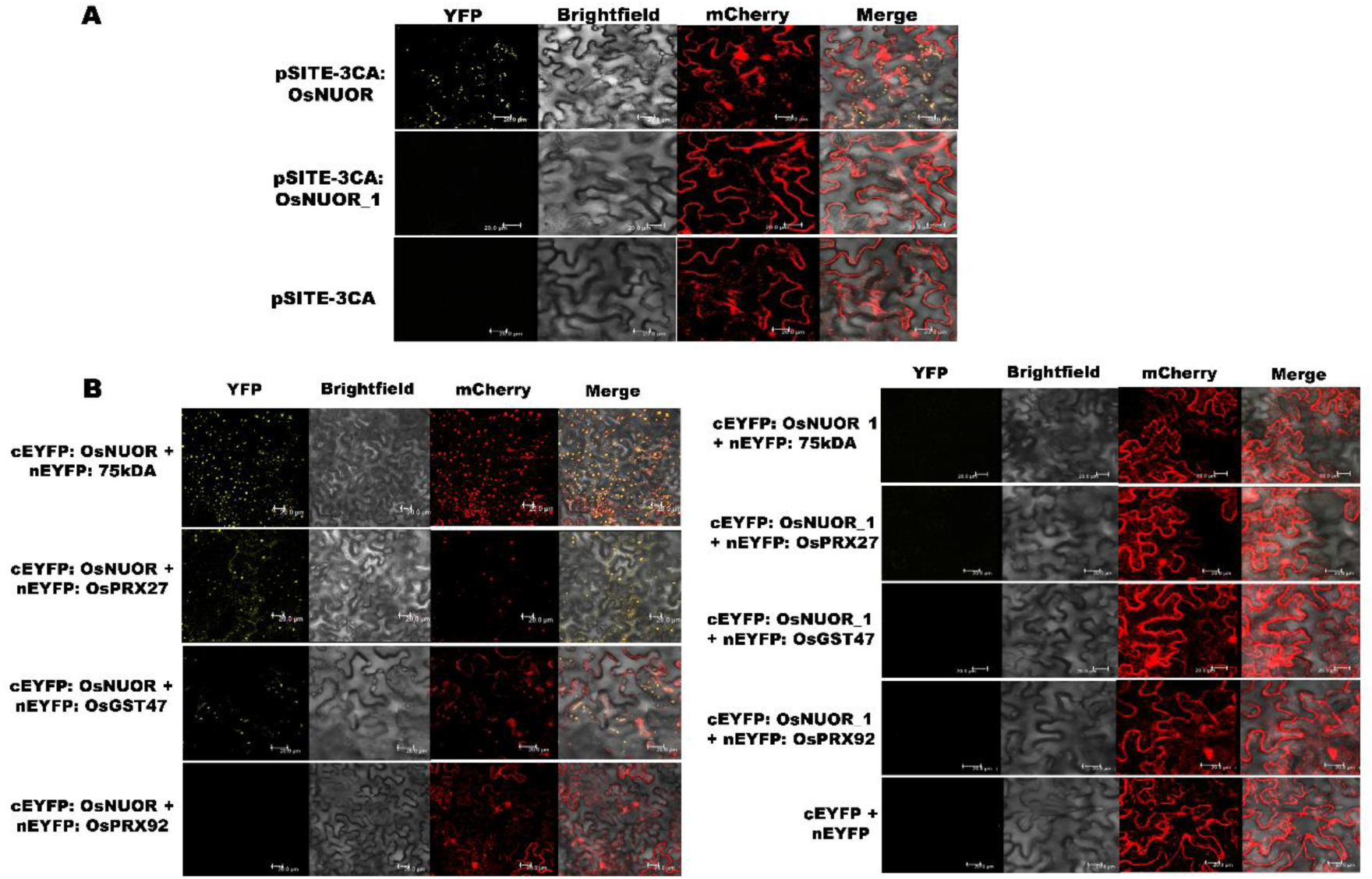
OsNUOR localizes in mitochondria and interacts with the predicted target proteins. **(A)** Confocal microscopic images showing co-colonization of YFP-tagged OsNUOR and mCherry-tagged mitochondrial marker (CD3-991) in *N. benthamiana* leaves. OsNUOR_1, a OsNUOR variant (OsNUOR^Δ1-108aa^), fails to localize in mitochondria. The YFP signal was visualized under YFP (yellow fluorescent protein) filter, while mCherry was visualized under RFP (red fluorescent protein) filter. Scale bars = 20 μm. **(B)** BiFC assay showing the interaction of OsNUOR with the predicted target proteins (75 kDa subunit of complex-1, Prx27 and GST47) in *N. benthamiana* leaves. OsNUOR failed to interact with Prx92. OsNUOR_1 was unable to interact with the target proteins. The OsNUOR/OsNUOR_1 was expressed as a C-terminal YFP fusion (cYFP) protein, whereas the target proteins (75 kDa subunit of complex-1, Prx27, GST47, and Prx92) were co-expressed as N-terminal YFP fusion (nYFP) proteins. Images were taken under confocal microscopy and yellow fluorescence dots merging with the red fluorescence dots of CD3-991, reflect physical interaction in the mitochondria. Scale bars = 20 µm.

STRING analysis (https://string-db.org/ version 11.5) predicted interacting partners of *OsNUOR,* including 75-kDa subunit of complex 1, various class III peroxidase (Prx) and glutathione S-transferase (GST) proteins (Supplementary Figures 6A and 6B). Most of the interacting partners were transcriptionally induced upon *R. solani* infection in rice (Supplementary Figure 7). The bimolecular fluorescence complementation (BiFC) (Figure 2B) as well as yeast 2-hybrid (Y2H) (Supplementary Figure 8) assays revealed that OsNUOR physically interacts with 75 kDa subunit, Prx27 as well as GST47, but not with Prx92 protein. However, OsNUOR_1 failed to interact with the target proteins (Figure 2B, Supplementary Figure 8).

Computational analysis predicted that 75 kDa subunit is localized in mitochondria, while Prx27 and Prx92 are secreted, and GST47 is a cytoplasmic protein (Supplementary Table 1). The confocal microscopic analysis reflected that 75 kDa subunit, Prx27 and Prx92 proteins but not GST47 protein, localized in mitochondria (Supplementary Figure 9).

### *OsNUOR* is important for enhanced ROS accumulation and lipid peroxidation in rice upon *R. solani* infection

DCFDA (2’,7’-dichlorofluorescin diacetate) staining revealed that ROS accumulation was marginally increased in OE lines under uninfected conditions, and it was several folds enhanced under infected conditions, compared to WT/EV lines (Figure 3A). On the other hand, edited lines had reduced ROS accumulation, even under infected conditions. Several of the respiratory burst oxidase homologs (*RBOH*s) including *RBOH-D* (Figure 3B, Supplementary Figure 10) that encode NADPH oxidases, pivotal players in apoplastic ROS production ^18^, were upregulated in OE lines under infected conditions. In addition, expression of the *OsNADP-ME2* gene, encoding NADP malic enzyme 2 that is positively associated with RBOH-mediated ROS production ^19^ was enhanced in OE lines (Figure 3C). However, the edited lines had basal-level expression of *RBOH* homologs as well as *OsNADP-ME2*, even under infected conditions (Figures 3B and 3C; Supplementary Figure 10). It was noteworthy that the expression of antioxidative enzymes including *OsCAT* (catalase), *OsAPX* (ascorbate peroxidase), *OsMnSOD* (superoxide dismutase), and *OsGR* (glutathione reductase) was downregulated in the OE lines, but enhanced in the edited lines under infected conditions (Supplementary Figure 11). However, the expression of these antioxidant genes remained comparable in different rice lines, under uninfected conditions.

**Figure 3:**
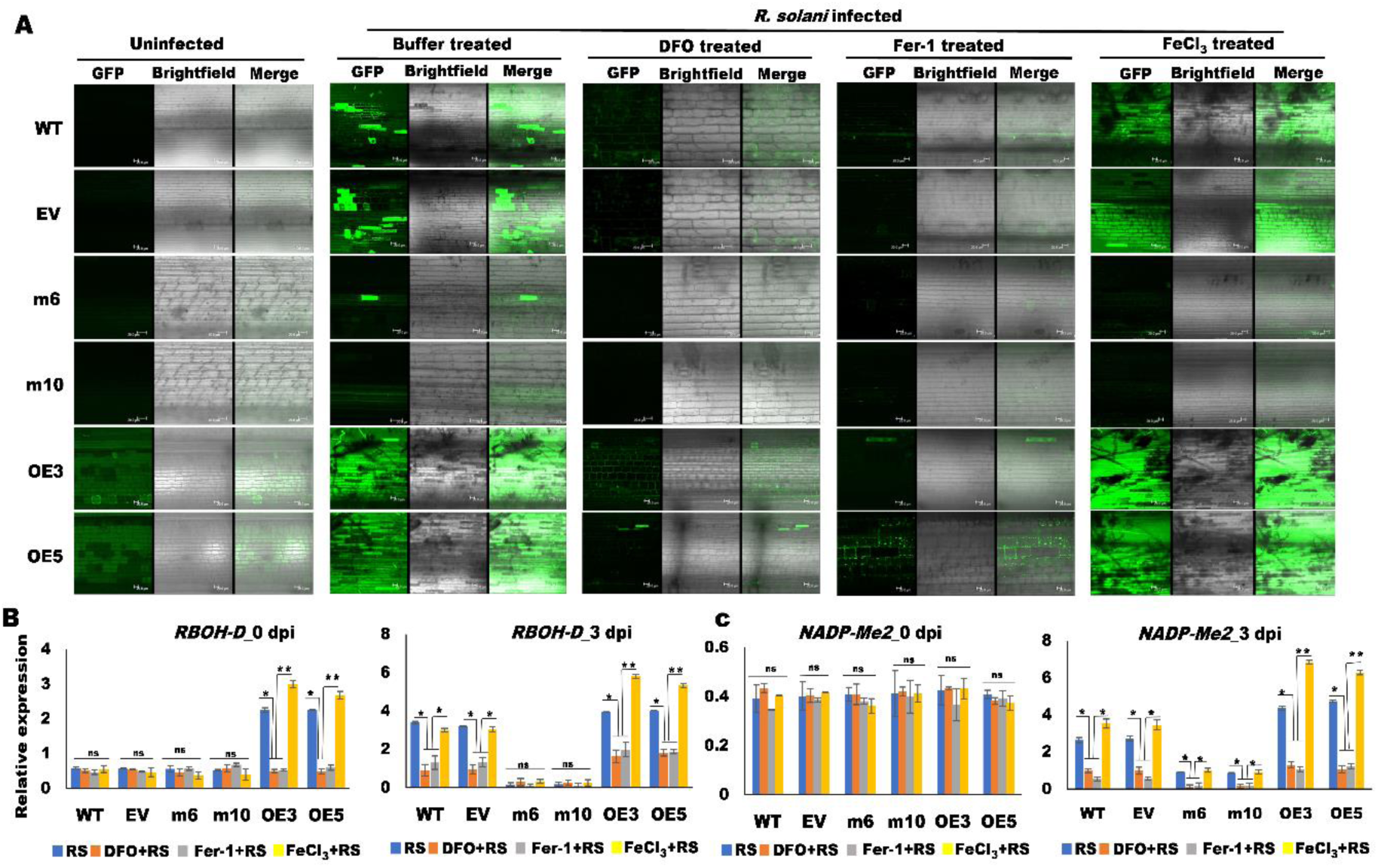
OsNUOR is important for enhanced ROS accumulation in rice upon *R. solani* infection. **(A)** DCFDA staining showing ROS accumulation (green fluorescing cells) in different rice lines, with and without *R. solani* infection. The effect of deferoxamine (DFO; 3mM), ferrostatin-1 (Fer-1; 10µM), and ferric chloride (FeCl_3_; 20 µM) treatment on ROS accumulation was also investigated. The images were recorded under GFP filter of Confocal Laser Scanning Microscope, using 20X objective. Scale bars = 20 µm. The relative expression of **(B)** Respiratory burst oxidase homolog D (*RBOH-D*) and **(C)** NADP malic enzyme 2 (*OsNADP-ME*2), at 0 dpi and 3 dpi. Relative expression was quantified using *actin* gene of *Oryza sativa* as an endogenous control. The average of three independent biological replicates is plotted and error bar represents mean ± standard error. ‘*’ and ‘**’ indicate significant differences at *p* < 0.05 and *p* < 0.001, respectively (estimated using one-way ANOVA and Student-Newman-Keuls test). ‘ns’ indicates no significant difference.

ROS burst causes peroxidation of unsaturated fatty acids and malondialdehyde (MDA) is used as a marker for phospholipid peroxidation in plants ^20^. We observed that under uninfected conditions MDA content was comparable in different rice lines, however, under infected conditions, it was significantly higher in OE lines but reduced in the edited lines, compared to WT/ EV plants (Figure 4A).

**Figure 4:**
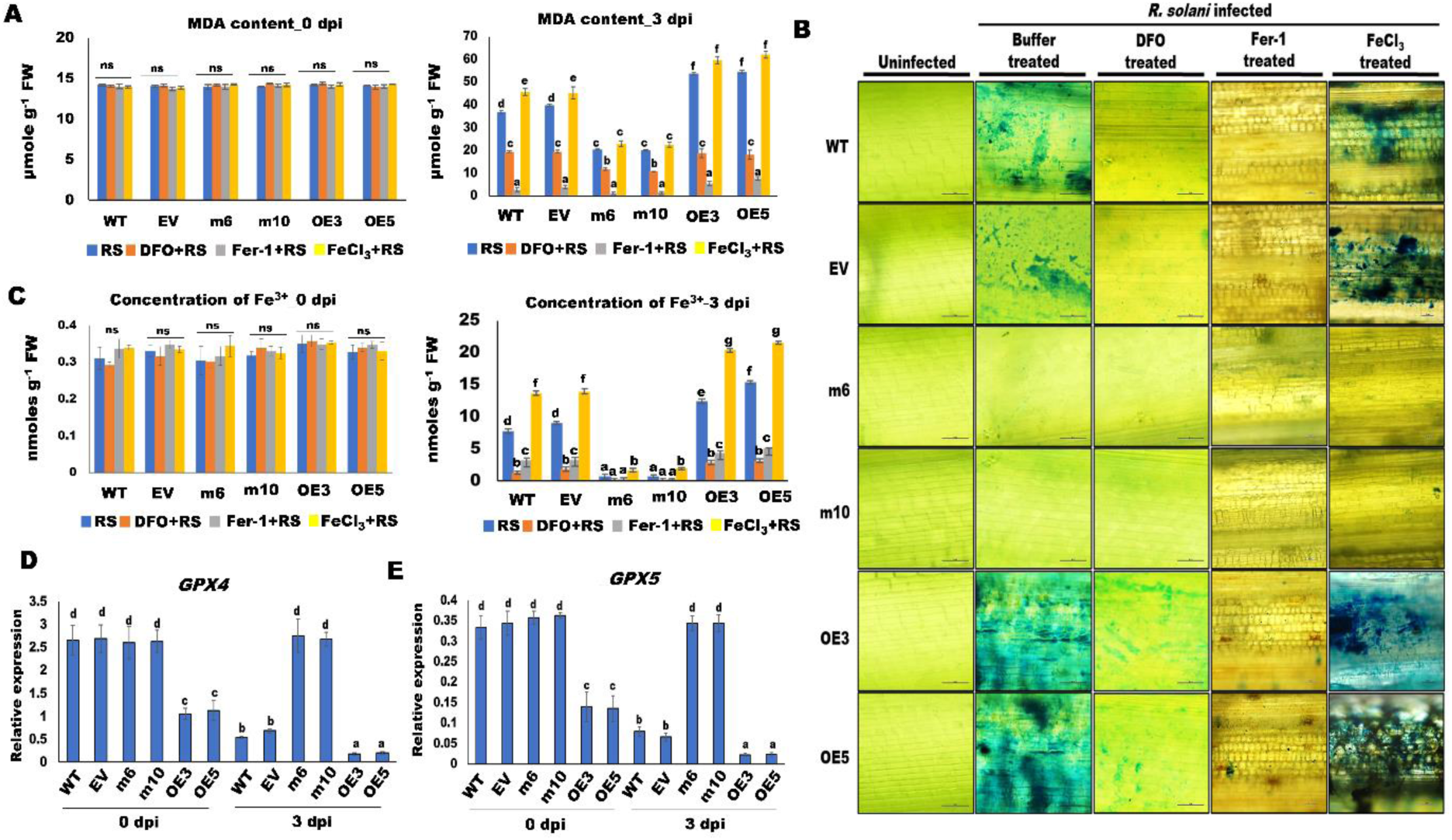
OsNUOR is important for enhanced Fe^3+^ accumulation and lipid peroxidation in rice under *R. solani* infected conditions. **(A)** Quantification of malondialdehyde (MDA) content, a lipid peroxidation marker, in different rice lines, at 0 and 3 dpi of *R. solani* infection. The effect of DFO (Deferoxamine; 3mM), Fer-1 (Ferrostatin-1; 10µM), and FeCl_3_ (Ferric chloride; 20 µM) treatment was also investigated. **(B)** Prussian blue staining (blue color) showing the accumulation of ferric ions (Fe^3+^) in different rice lines, at 0 and 3 dpi. Images were recorded under a Nikon 80i-epi-fluorescence microscope. Scale bars = 50 µm. **(C)** Quantification of ferric ion (Fe^3+^) in different lines at 0 and 3 dpi. Relative expression of **(D)** *Glutathione peroxidase 4* (*OsGPX4*) and **(E)** *Glutathione peroxidase 5* (*OsGPX5*) genes, at 0 and 3 dpi. Relative expression was quantified using *actin* gene of *Oryza sativa* as an endogenous control. The average of three independent biological replicates is plotted and the error bar represents mean ± standard error. Different letters represent a significant difference at *p* < 0.05 (one-way ANOVA, Student-Newman-Keuls test).

### *OsNUOR* is required for enhanced Fe^3+^ accumulation and cell death in the infected tissues

Considering that an enhanced level of ferric iron (Fe^3+^) is associated with lipid peroxidation ^21,22^, we analyzed Fe^3+^ accumulation in rice using Prussian blue staining ^23,24^. Under infected conditions, staining was intense in OE lines, whereas edited lines showed limited staining, comparable to the uninfected controls (Figure 4B). Moreover, Fe^3+^ content (quantified using a commercially available kit) was enhanced in OE lines, but reduced in the edited lines, compared to WT and EV lines (Figure 4C). Under uninfected conditions, Fe^3+^ content was comparable in different rice lines.

To investigate the importance of Fe^3+^ accumulation and lipid peroxidation during *R. solani* infection, we treated rice with deferoxamine (DFO), an iron chelator ^25^, ferrostatin-1 (Fer-1), a lipid peroxidation inhibitor ^26^, and FeCl_3_, as a source of Fe^3+^ ion ^27^. Notably, ROS accumulation (Figure 3A), MDA (Figure 4A), and Fe^3+^ (Figures 4B and 4C) content were enhanced by FeCl_3_, but reduced by DFO/Fer-1 in *R. solani*-infected WT/EV/OE lines. In addition, the expression of *RBOH* homologs (Figure 3B; Supplementary Figure 10) and *OsNADP-ME2* (Figure 3C) was significantly downregulated by DFO/Fer-1, whereas FeCl_3_ treatment enhanced their expression in WT/EV/OE lines. DFO/Fer-1/FeCl_3_ treatment failed to alter the expression of *RBOH* homologs and *OsNADP-ME2* in the edited lines. Notably, disease symptoms (Figure 5A), RVSC index (Figure 5B), pathogen load (Figure 5C), as well as fungal infestation (Figure 5D) were enhanced in FeCl_3_ but suppressed in DFO/Fer-1 treated rice lines. On the other hand, FeCl_3_ treatment failed to impart disease susceptibility in the edited lines, and MDA content (Figure 4A), Fe^3+^ content (Figure 4C), as well as ROS accumulation (Figure 3A) remained unaltered.

**Figure 5:**
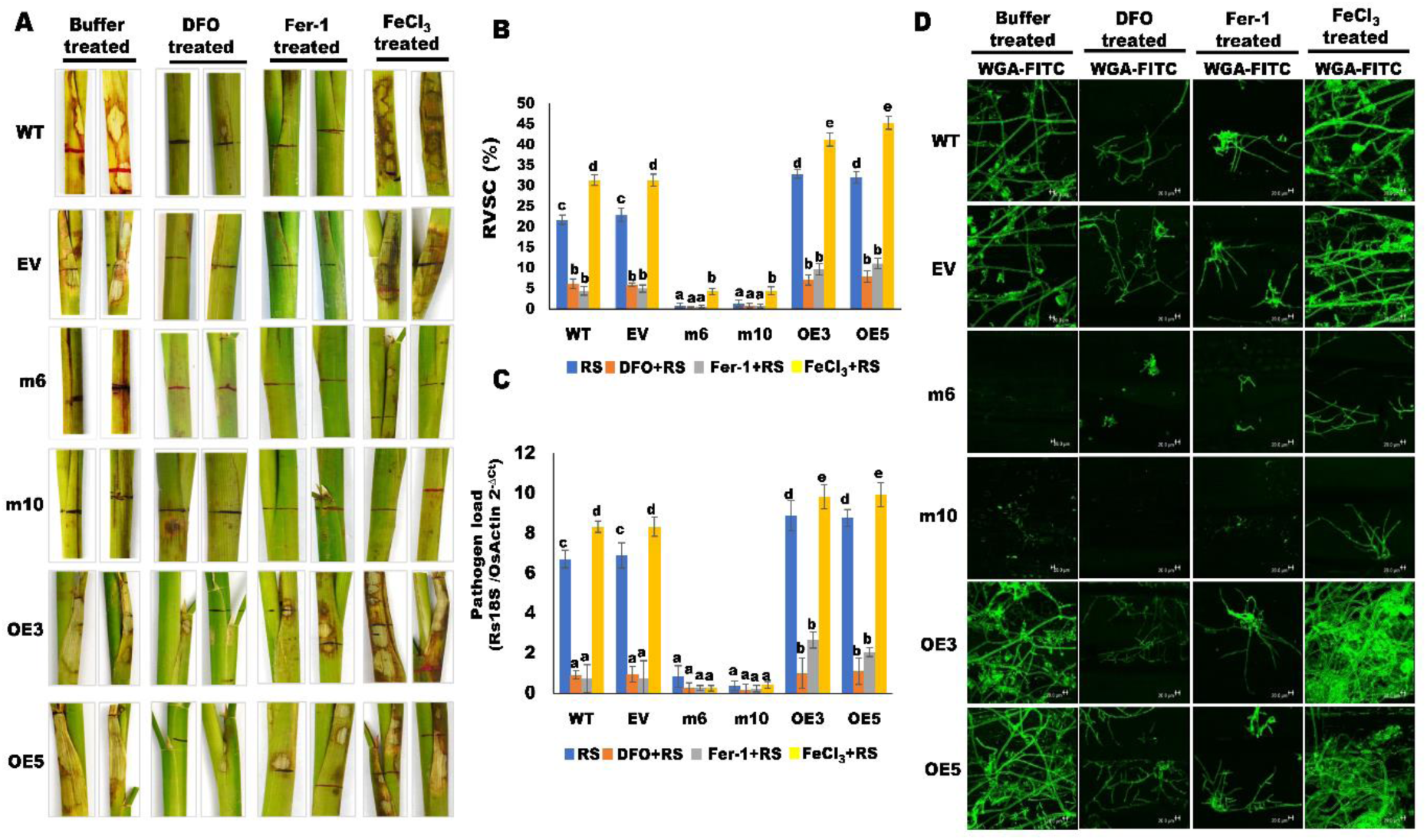
Deferoxamine and ferrostatin-1 treatment imparts disease tolerance whereas FeCl_3_ treatment enhances disease susceptibility. **(A)** Disease symptoms in different rice lines upon *R. solani* infection, at 5 dpi. The effect of Deferoxamine (DFO; 3mM), ferrostatin-1 (Fer-1; 10µM), and ferric chloride (FeCl_3_; 20 µM) treatment on *R. solani* infection are depicted. **(B)** RVSC (relative vertical sheath colonization) based disease index and **(C)** Bar graph showing qRT-PCR-based quantification of *R. solani* 18S rRNA gene (as pathogen load) in different lines. The relative expression was measured using ΔCt method, where ΔCt is the difference between Ct value of *18S rRNA* of *R. solani* and *Oryza sativa actin* gene. **(D)** WGA-FITC staining reflects *R. solani* mycelial growth in different lines. The mycelia were observed under GFP filter of the Confocal Laser Scanning Microscope AOBS TCS-SP5, Leica, Wetzlar, Germany) using 20X objective. Scale bars = 20 µm. The bar denotes the average of three independent biological replicates and error bar represents mean ± standard error. Different letters represent a significant difference between samples at *p* < 0.05 (one-way ANOVA, Student-Newman-Keuls test).

The above analyses emphasized that the *R. solani* infection induces an iron-dependent host cell death and OsNUOR is an important player in its elicitation. Iron-dependent cell death is referred as ferroptosis, characterized by enhanced accumulation of lipid ROS ^28,29^. One of the hallmarks of ferroptosis is the downregulation of glutathione peroxidase 4 (*GPX4*), a phospholipid hydroperoxidase that prevents uncontrolled lipid peroxidation ^30–32^. We observed that *GPX4* homologs in rice (*OsGPX4* and *OsGPX5*) were significantly downregulated under *R. solani*-infected conditions in WT/EV/OE lines, whereas they remained unaltered in the edited lines (Figures 4D and 4E).

### Redox status of the infected tissues is important for *OsNUOR* expression

We observed that the expression of *OsNUOR* was enhanced under infected conditions, repressed by DFO/Fer-1, and induced by FeCl_3_ treatment in WT/EV/OE lines (Figure 6A). On the other hand, edited lines had significantly reduced *OsNUOR* expression, even under infected conditions, with and without DFO/Fer-1/FeCl_3_ treatment. Considering the above, we anticipated that redox signaling may modulate *OsNUOR* expression under *R. solani-*infected conditions. This was reinforced by reporter GUS assay, wherein *GUS* expression under *OsNUOR* promoter was induced upon *R. solani* infection (Supplementary Figures 12A-C). The DFO/Fer-1 treatment suppressed, whereas FeCl_3_ treatment enhanced GUS expression (Figures 6B and 6C). Notably, FeCl_3_ treatment induced GUS expression, even under uninfected conditions. Similarly, qRT-PCR analysis reflected that DFO/Fer-1 suppresses, whereas FeCl_3_ enhances *OsNUOR* expression in rice (Figure 6D).

**Figure 6:**
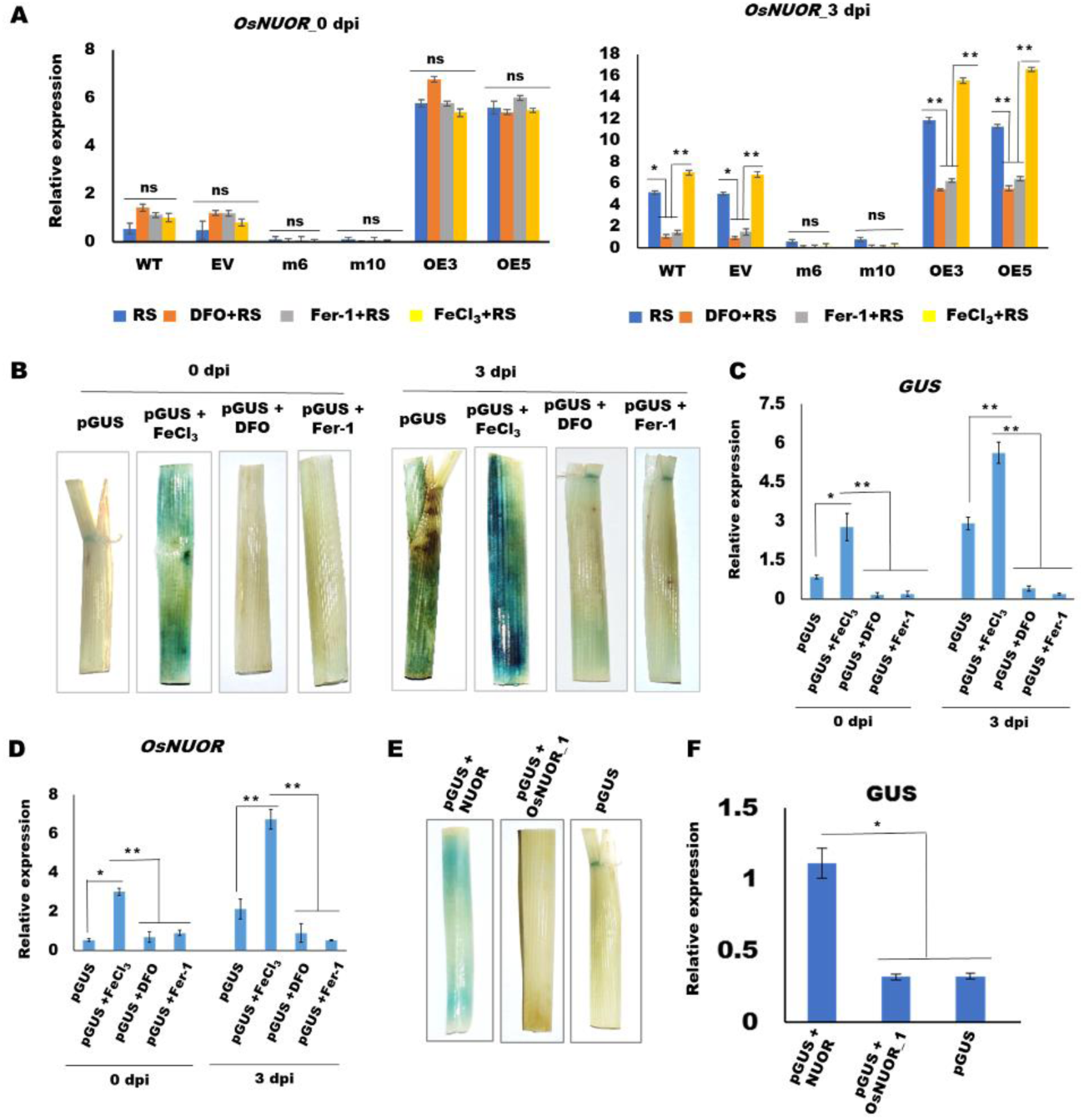
Redox status of *R. solani*-infected rice tissues regulates *OsNUOR* expression. **(A)** The relative expression of *OsNUOR* in different rice lines. The effect of deferoxamine (DFO; 3mM), ferrostatin-1 (Fer-1; 10µM), and ferric chloride (FeCl_3_; 20 µM) treatment on gene expression is shown. **(B)** GUS reporter assay, showing the effect of DFO/Fer-1/FeCl_3_ treatment and *R. solani* (RS) infection on activation of *OsNUOR* promoter (reflected as the appearance of blue colour). Relative expression of **(C)** β-glucuronidase (GUS) and **(D)** *OsNUOR* in rice sheaths, quantified by qRT-PCR using rice *actin* gene as endogenous control. **(E)** GUS reporter assay reflecting the induction of OsNUOR promoter upon co-expression of *OsNUOR* in rice. ‘pGUS’ represents leaves infiltrated with *OsNUOR* promoter construct. **(F)** qRT-PCR-based quantification of *β-glucuronidase* (GUS) expression, using rice *actin* gene as endogenous control. The average of three independent biological replicates is plotted and error bar represents mean ± standard error. ‘*’ and ‘**’ indicate significant differences at *p* < 0.05 and *p* < 0.001, respectively (estimated using one-way ANOVA and Student-Newman-Keuls test). ns indicates no significant difference.

We observed that agrobacterium-mediated transient overexpression of OsNUOR, but not OsNUOR_1 enhances GUS expression under *OsNUOR* promoter, in rice (Figures 6E and 6F). This tempted us to speculate that a cross-talk between OsNUOR and ROS signaling modulates gene expression and sheath blight disease susceptibility in rice. The alteration of the cross-talk as observed in the edited lines imparts disease tolerance in rice.

## Discussion

Imparting sheath blight disease (SBD) resistance remains challenging for sustainable rice cultivation. Being a polyphagous necrotrophic fungal pathogen, a practical level of disease resistance remains elusive in the available rice germplasm. Even though rice growers spend millions of dollars to mitigate SBD, huge annual yield loss is reported due to the disease ^33^. It is hypothesized that *R. solani* infection modulates the host’s physiological and metabolic functions that render disease susceptibility in rice ^4,6,,14^. Here, we provide evidence that an alternative NADH:ubiquinone oxidoreductase (OsNUOR) functions as a susceptibility factor for SBD. We demonstrate that overexpression of *OsNUOR* enhances disease susceptibility in rice, whereas knock-down of the gene (by Cas9-based genome editing) imparts disease resistance. Notably, edited lines did not have any phenotypic alteration, and due to the transgene-free nature, they are subject to lesser biosafety regulations before deployment under farmer’s fields. Hence, our study reports an effective biotechnological intervention to mitigate SBD, the importance of which has been highlighted in an economic surveillance report, that by the cultivation of disease-resistant cultivars (once available), the United States can have surplus rice to feed 1.7 million people with an average consumption of 65 kg/person/year ^33^. The implication will be several-fold higher if adopted worldwide.

Previous studies have shown that the necrotrophic phase of *R. solani* infection is associated with enhanced ROS accumulation and cell death in rice ^4,14^. Here, we observed that compared to the WT/EV lines, ROS accumulation was enhanced in *OsNUOR* overexpression (OE) lines, but reduced in the edited lines, under infected conditions. Notably OE lines had marginally enhanced ROS levels, even under uninfected conditions. Various studies have demonstrated that members of alternative NUOR, due to NADH dehydrogenase activity, participate in the electron transport chain (ETC) and promote respiration ^11,13,,34^. We anticipate that via interaction with 75 kDa subunit of complex-1, OsNUOR potentially stimulates ETC, and electron leakage from the ETC contributes to cellular ROS production ^35^. Moreover, via interaction with Prx proteins that can generate hydroxyl radicals (**·**OH) in a Fenton-type reaction ^36,37^, OsNUOR may participate in ROS production.

In general, plants upregulate their antioxidant machinery to keep ROS under a threshold ^38,39^, however, we observed that genes encoding antioxidant enzymes, including Catalase (*OsCAT*), Ascorbate peroxidase (*OsAPX*), Manganese superoxide dismutase (*OsMnSOD*), and Glutathione reductase (*OsGR*) were downregulated in WT, EV as well as OE lines, while they were induced in edited lines under *R. solani*-infected conditions. These results suggest that OsNUOR has double-edged functions, it enhances ROS production and also impairs the antioxidative defense system in the infected tissues. It is possible that via physical interactions, OsNUOR sequesters GSTs in mitochondria, and alters their sub-cellular functions, thereby impairing the antioxidant defense system. The role of glutathione S-transferases (GSTs) in ROS scavenging ^40^ and SBD tolerance ^41^ has been previously reported. It is also possible that OsNUOR-mediated ROS production may modulate transcriptional machinery to create an oxidative stress-enriched environment in rice. Indeed, we observed that modulation of the redox status of the infected cells by DFO/Fer-1 (which inhibited ROS accumulation) suppressed, whereas FeCl_3_ (which enhanced ROS accumulation) induced the expression of not only *OsNUOR* but also *OsNADP-Me2* (NADP malic enzyme 2) and *RBOH* homologs (NADPH oxidases) in the OE lines, but not in the edited lines. It is to be noted NADP malic enzymes supply electrons to cytosolic electron donor NADPH that further reduces RBOH proteins to promote ROS generation ^23,42^.

An enriched redox environment causes peroxidation of polyunsaturated fatty acids which triggers a chain of events to damage cellular/organellar membranes ^43–45^. We observed MDA content (a marker for lipid peroxidation ^46,47^ was enhanced in OE lines but not in the edited lines. Similarly, ferric ion accumulation was enhanced in the infected OE lines, but reduced in the edited lines, as revealed by Prussian blue staining, as well as biochemical estimation. Recent studies have shown that lipid ROS and accumulated ferric ions induce a specialized form of iron-dependent cell death, known as ferroptosis ^21,48^. The impairment of antioxidative systems, particularly the glutathione-dependent system and/or production of oxygen-derived free radicals from the Fenton reaction (Fe^2+^ + H_2_O_2_ → Fe^3+^ + HO· + HO), are known to trigger ferroptosis ^48–50^. We observed that rice homologs of glutathione peroxidase 4 (*OsGPX4* and *OsGPX5*) were significantly downregulated in the WT/EV/OE lines but not in edited lines, under *R. sola*ni-infected conditions. Moreover, we observed that treatment with Fer-1/ DFO, the known inhibitors of ferroptosis, not only prevented ROS accumulation, lipid peroxidation, and ferric ion accumulation but also suppressed SBD in WT/EV/OE rice. On the other hand, exogenous treatment with ferric chloride enhanced disease susceptibility along with induced accumulation of lipid ROS in the WT/EV/OE lines, but not in the edited lines. Considering the above, we propose that the necrotic disease symptoms observed in *R. solani*-infected tissues are due to ferroptosis and OsNUOR is an important player in its elicitation.

Overall, our study emphasizes that *R. solani* infection creates an oxidative stress-enriched environment that upregulates *OsNUOR*, which contributes to enhanced ROS and ferric ions accumulation, together they induce iron-dependent lipid peroxidation and cell death. The edited lines are defective in these processes and therefore exhibit a high level of disease tolerance (Figure 7). Further research is important to investigate OsNUOR-mediated redox signaling in plants and identify newer players of ferroptosis that can be modulated for imparting sheath blight disease tolerance in rice.

**Figure 7:**
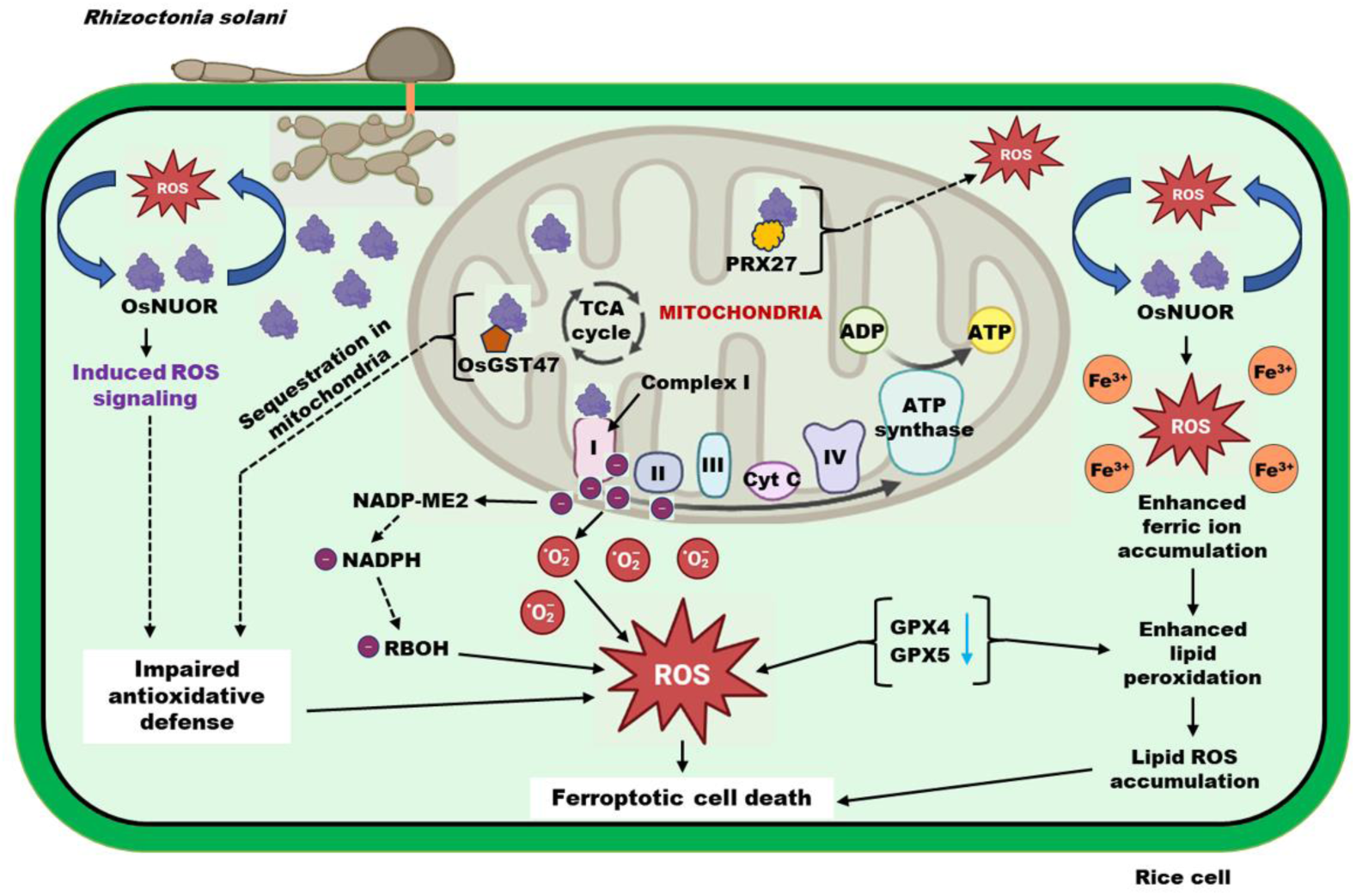
Proposed role of OsNUOR in promoting sheath blight disease susceptibility in rice by inducing ROS signaling and ferroptosis. *R. solani* infection creates an oxidative stress-enriched environment, which upregulates *OsNUOR* expression. OsNUOR gets localized into mitochondria, interacts with the 75 kDa subunit of complex-I, and potentially stimulates the electron transport chain (ETC). The electron leakage from the ETC converts molecular oxygen to superoxides (O_2_^•−^) and other reactive oxygen species (ROS). The OsNUOR also interacts with Prx27 and contributes to ROS production. Further, enhanced NADP-malic enzyme (NADP-ME2) and respiratory burst oxidases (RBOHs) contribute to enhanced ROS production. In addition, OsNUOR potentially sequesters OsGST47 in the mitochondria and prevent its sub-cellular functions, overall impairing the antioxidative defense and enhancing ROS accumulation. The ROS signaling causes transcriptional changes, leading to reduced expression of antioxidative enzymes, including catalase, glutathione reductase, ascorbate peroxidase, and glutathione peroxidases (GPX4 and GPX5). The ROS along with enhanced accumulation of ferric ions contribute to lipid peroxidation. Altogether, the enhanced accumulation of ferric ions and lipid ROS, along with depletion of GPX and impaired antioxidative defense induces ferroptosis, leading to necrotic cell death in rice upon *R. solani* infection. However, in *OsNUOR* edited lines accumulation of ferric ions and lipid ROS were compromised, whereas the expression of GPXs (GPX4 and GPX5) and various antioxidative enzymes were enhanced. Therefore, ferroptosis was repressed and disease resistance was observed in the edited lines.

## Materials and methods

### Biological materials

Rice, *Oryza sativa* ssp. *japonica* (cv. TP309) (wild-type; WT) was used in the study. The plants were grown in pots under greenhouse conditions (28°C temperature, 80% relative humidity, 16/8 h of day/night cycle, and 250 µmol light intensity) ^6^. The tobacco (*Nicotiana benthamiana*) plants were grown in soilrite (Keltech Energies Ltd, India) in a growth chamber (Conviron, USA at 26°C under 12/12 h of day/night cycle and 70% relative humidity ^51^. The fungal pathogen *Rhizoctonia solani* AG1-IA strain BRS1 ^52,53^ was grown on PDA (Potato-dextrose agar; Himedia, Mumbai, India; 39 g/L) plates at 28°C, as described earlier ^54^. *Escherichia coli* (DH5α) was grown on LB (Luria bertani) media with required antibiotic selection at 37°C. Similarly, *Agrobacterium tumefaciens* strains (EHA105- used for transformation in rice; GV3101- used for transformation in *Nicotiana benthamiana*) were grown in Yeast Extract Peptone (YEP; Himedia) media at 28°C. Yeast cells were grown on YEPD (yeast extract peptone dextrose) media at 30°C (MP Biomedicals) as described earlier ^55^.

### In-silico analysis

The gene coding sequence (CDS) was retrieved from the Rice Genome Annotation Project (RGAP) database (https://rice.uga.edu/). The promoter sequence was retrieved from PLAZA 4.0 database (https://bioinformatics.psb.ugent.be/plaza). The STRING Protein-Protein Interaction Networks (https://version-11-5.string-db.org/) was used to predict potential interacting partners. The signal peptide and subcellular location of proteins were predicted by SignalP-5.0 (https://services.healthtech.dtu.dk/services/SignalP-5.0/) and TargetP-2.0 (https://services.healthtech.dtu.dk/services/TargetP-2.0/), respectively. The interpro scan (https://www.ebi.ac.uk/interpro/search/sequence/) was used to predict the conserved domain.

### Generation of NUOR overexpressing rice lines

The coding sequence of Os*NUOR* (1689 bp) was PCR amplified from rice (cv. TP309) cDNA using OsNUOR-FP and OsNUOR-RP primer pair and cloned into pMDC84 (ABRC stock number: CD3-743) vector to generate an overexpression construct (CaMV35S:OsNUOR:GFP). The construct was transformed into rice calli (cv. TP309) using *Agrobacterium-*mediated transformation to raise overexpression (OE) lines, as described previously ^56,57^. The CaMV35S-FP/ OsNUOR-RP primers were used to confirm the presence of transgene. Similarly, the presence of T-DNA was confirmed by PCR using HPT-FP/HPT-RP primer pairs (Supplementary Figure 1). Each of these lines was propagated to T_2_ generation and two independent OE lines having higher fold expression of *OsNUOR* gene were selected for further analysis. List of primers used during the study is listed in (Supplementary Table 2).

### CRISPR-Cas9 mediated generation of OsNUOR-edited rice lines

For Cas-9 mediated genome editing of *OsNUOR* (LOC_Os07g37730), two guide RNA having minimum off-targets and maximum on-target scores were selected using CRISPR-P 2.0 tool (http://crispr.hzau.edu.cn/CRISPR2/). Primers were designed corresponding to guide RNA target to amplify polycistronic transfer RNA-gRNA-scaffold (sgRNA) from template plasmid pGTR (Addgene plasmid #63143). Using Golden Gate assembly kit [(NEBridge® Golden Gate Assembly Kit (BsaI-HF® v2)], sgRNAs were assembled and amplified as a single transcript, as described before ^58^. The amplified construct was cloned into pRGEB32 (Addgene plasmid #63142) vector under rice ubiquitin promoter (*OsUbi*) for Cas-9 and rice snoRNA U3 promoter for guide-RNA, using Golden Gate assembly kit. The positive construct was mobilized in *A. tumefaciens* EHA105 and the recombinant strain was used for the transformation of 25-day-old rice (cv. TP309) callus, as described in ^57,59^. The edited plants were screened using primer pair HPT-FP/HPT-RP and Cas9-FP/Cas9-RP followed by Sanger DNA sequencing. Each of these lines were propagated to T_2_ generation and two independent homozygous edited lines were used for further characterization. List of primers used during the study is listed in (Supplementary Table 2).

### *R. solani* infection assays

The sclerotia of *Rhizoctonia solani* AG1-IA isolate (BRS1) were used to infect tillers of 45-day-old rice lines, as described before ^52^. The infected plants were incubated in a growth chamber under high humidity and monitored for the appearance of disease symptoms. The relative vertical sheath colonization (RVSC) was calculated as described in ^52^. At least, 20 tillers of each line were analyzed in each experiment and experiments were independently repeated three times. The infected tillers were photographed using a DSLR camera at 5 dpi.

### WGA-FITC staining

The colonization of *R. solani* in the rice sheaths was visualized using confocal microscopy. For this, freshly harvested sheaths were boiled in 1 M KOH (Amresco Inc.) for 30 min and subsequently neutralized with 50 mM Tris (pH 7.0) followed by staining with wheat germ agglutinin (WGA)-FITC (Sigma-Aldrich) solution (20 µg/ml) for 30 min ^54,60^. Upon washing with sterile milliQ water, the samples were analysed under a GFP filter of Confocal Laser Scanning Microscope (AOBS TCS-SP5, Leica, Wetzlar, Germany). The images were processed using LAS AF Version: 2.6.0 build 7266 software (Leica). The experiment was performed in three biological and three technical replicates.

### Detection of ROS burst

ROS accumulation in the infected tillers was detected using ROS-sensitive dye H_2_DCFDA (2′,7′- dichlorodihydrofluorescein diacetate)^4^. For this, infected rice sheaths were excised and immersed in milliQ water for 5 min ^61^. Thereafter, the sheaths were stained with 10µM H_2_DCFDA (Thermo Fisher Scientific Inc. USA) solution prepared in phosphate-buffered saline (PBS) buffer (10 mM, pH 7.4) and incubated in a desiccator for 30 min. ROS accumulation was observed under the GFP filter of the Confocal Laser Scanning Microscope (AOBS TCS-SP5, Leica, Wetzlar, Germany). The images were analyzed using LAS AF Version: 2.6.0 build 7266 software (Leica). The experiment was performed in three biological and technical replicates.

### Detection of ferric iron accumulation by Prussian staining

To monitor ferric iron (Fe^3+^) accumulation, the rice sheaths were incubated in 7% (w/v) potassium ferrocyanide (Sigma-Aldrich) and 2% (v/v) hydrochloric acid (1:1, v/v) for 15 h at room temperature, as described earlier ^62^. The Fe^3+^ reacts with ferrocyanides to form a bright blue pigment called Prussian blue or ferric ferrocyanides. The stained sheaths were observed under a Nikon 80i-epi-fluorescence microscope (Eclipse Ni-E, NCY-NI-E-931446, Japan). The images were recorded and analyzed using NIS-elements AR 5.41.00 imaging software. The experiment was performed using three biological and technical replicates.

### Iron quantification

0.5 g of the rice tissues (5-6 excised sheaths pooled) were homogenized in 1 ml of 80% acetone using liquid nitrogen. After 30 min incubation, the homogenate was centrifuged at 10000 rpm for 10 min and the supernatant was collected for iron quantification using an Iron Assay kit (ab83366; Abcam, Cambridge, UK), as per manufacturer’s protocol. Briefly, the samples, with (for estimation of total iron) and without (for Fe^2+^ estimation) iron reducer, were incubated with an iron probe in the dark for 30 min, and absorbance (OD) at 593 nm was measured using Multiskan SkyHigh Microplate Spectrophotometer (Thermo Scientific, USA). The total iron/Fe^2+^ ion content was estimated using a standard curve, and Fe^3+^ content was calculated by deducting Fe^2+^ content from the total iron content. The experiment was performed using three biological and three technical replicates.

### Malondialdehyde (MDA) estimation

0.5 g rice tissues (5-6 excised sheaths pooled) were homogenized in 1 ml of 80% acetone, and incubated on ice for 30 min. Upon centrifugation at 10000 rpm for 10 min, the supernatant was collected for quantification of MDA, using a thiobarbituric acid reactive substances (TBARS) Assay Kit (700870; Cayman Chemical, Ann Arbor, MI, USA), following manufacturer’s protocol. The samples were mixed in 1:1 ratio with TCA Assay Reagent (10%) followed by the addition color reagent and boiling in a water bath for 1h. Subsequently, samples were centrifuged at 8000 rpm for 10 min, supernatant was collected and after 30 min of incubation absorbance (OD) at 535 nm was measured using Multiskan SkyHigh Microplate Spectrophotometer (Thermo Scientific, USA). The standard curve for MDA was used to extrapolate MDA content in the respective samples. The experiment was performed in three independent biological replicates and three technical replicates.

### Deferoxamine, Ferrostatin-1 and Ferric chloride treatment

Deferoxamine (DFO, 3mM, Sigma-Aldrich), Ferrostatin-1 (Fer-1, 10µM, Sigma-Aldrich), and Ferric chloride (FeCl_3_, 20 µM, Sigma-Aldrich) solutions were prepared in a sterile MilliQ. The chemical solutions were sprayed onto 45-day-old tillers of rice plants, and after 24 h, the plants were infected with *R. solani*. The disease occurrence was recorded and samples were harvested for further analysis.

### GUS reporter assay

The promoter region (1000 bp) of the *OsNUOR* gene was cloned in pGWB3 vector ^63^, being transcriptionally fused with *uidA* (β-glucuronidase) reporter gene (pGWB3:*OsNUOR:Promoter*-GUS) and mobilized into *Agrobacterium* strain (EHA105). The recombinant strain was infiltrated into the midvein of 45-day-old rice plants using a syringe, as described earlier ^64^, followed by the spray of DFO/Fer-1/FeCl_3_ solutions, the next day. After 24 h, the tillers were infected with *R. solani* and incubated under controlled conditions. After 3 dpi, the tillers were harvested and incubated in X-gluc solution for GUS assay, as described earlier in^55^. After overnight incubation at 37°C, samples were destained in an ethanol and acetic acid (3:1) solution. The destained sheaths were photographed and images were captured using a DSLR camera. Expression of the *GUS* (*β-glucuronidase*) gene was quantified by qRT-PCR using a gene-specific primer pair (Supplementary Table 2). The experiment was independently repeated at least three times and in each experiment a minimum of 10 tillers/samples were analyzed.

### cDNA synthesis and expression analysis

The 5-6 excised sheaths were pooled and used for total RNA isolation using a RNeasy Plant RNA isolation kit (Qiagen, Valencia, CA). 1µg of total RNA was used to synthesize cDNA using Verso cDNA synthesis kit (Thermo Fisher Scientific Inc, USA). The expression of genes was quantified by qRT-PCR using gene-specific primers (Supplementary Table 2). Rice *actin* gene was used for the normalization of gene expression. The relative expression was estimated by the 2^−ΔCt^ method ^14^ and fold change was calculated by 2^-ΔΔCt^ method ^65,66^. The mean of three independent biological experiments was used to calculate the standard error.

### Cloning of OsNUOR variant

A variant of OsNUOR i.e OsNUOR-1 (OsNUOR**^Δ1-108^**) was created by deleting the 1-108 aa from N-terminus. For this, the 325-1686 bp region was PCR amplified from OsNUOR gene construct and cloned into pENTR/D-TOPO entry vector (pENTR / D-TOPO Cloning Kit, Thermo Fisher Scientific Inc, USA). Subsequently, this entry construct was used to mobilize into different destination vectors, including pSITE3CA; for localization study, pSITE-cEYFP-N1; for BiFC study, and pGADT7-AD; for Y_2_H study.

### Yeast two-hybrid assay

Full-length CDS of OsNUOR target proteins [75 kDA subunit, Prx92, Prx27, OsGST47] were PCR amplified from rice cDNA using gene-specific primer pairs (Supplementary Table 2) and cloned individually in the pGBKT7-BD bait vector (Clontech, USA). The *OsNUOR* and *OsNUOR_1* were cloned in pGADT7-AD prey vector (Clontech, USA). The bait plasmids were individually transformed into the Yeast two-hybrid (Y2H) Gold yeast strain as per manufacturer’s protocol (EZ-Yeast transformation kit, MP Biomedicals, USA). Subsequently, the prey construct was mobilized into the yeast strains expressing bait constructs. The positive transformants were selected on SD - Leu-Trp double dropout plates. The interaction was confirmed by growing transformed yeast cells on SD-Leu-Trp-Ade-His quadruple drop-out plates supplemented with 15 mM 3-Amino-1, 2, 4-triazole (3AT) incubated at 30°C for 3 days ^51^. The yeast cells co-transformed with pGADT7-P53 and pGBKT7-RecT were used as a positive control.

### Localization

The *OsNUOR* and *OsNUOR_1* were cloned into destination vector pSITE3CA (GenBank: EF212293.1) having C-terminal YFP fusion and transformed into *A. tumefaciens* GV3101. The YFP fusion constructs harbouring agrobacterial strain was mixed in 1 : 1 ratio with the strain expressing mCherry-tagged mitochondrial marker CD3-991 ^17^ in infiltration buffer as mentioned earlier by ^51^ and infiltrated into 3-weeks-old *N. benthamiana* leaves ^51^. After 2 days of infiltration, the YFP were visualized under a Leica TCS SP5 confocal scanning microscope (Leica Microsystems) with appropriate lasers [514/527 nm (excitation/emission) for YFP and 581/644 nm (excitation/emission) for mCherry].

### Bimolecular fluorescence complementation (BiFC) assay

OsNUOR/ OsNUOR_1 and different targets (75 kDa subunit, Prx92, Prx27, OsGST47) were cloned in pSITE-cEYFP-N1 (CD3-1651) and pSITE-nEYFP-C1 (CD3-1648) vectors, respectively. The constructs were individually transformed into *A. tumefaciens* GV3101 competent cells. The agrobacterial strain expressing *OsNUOR/OsNUOR_1* was mixed with the strain expressing the target proteins along with mCherry-tagged mitochondrial marker (CD3-991) ^17^ in infiltration buffer (100 mM MgCl_2_, 0.5 mM MES buffer, and 200 mM acetosyringone), incubated for 2 h, and co-infiltrated into the *N. benthamiana* leaves, as described in ^55^. After 2 d of infiltration, the reconstitution of YFP fluorescence was analyzed under a 514/527 nm (excitation/emission) and mCherry was analyzed under 581/644 nm (excitation/emission) in Leica TCS SP5 confocal scanning microscope (Leica Microsystems).

### Phenotypic analysis of rice lines

The plant height (65-day-old, average of 20 plants, in cm), panicle length (average of 20 panicles, in cm), seed weight (average of 100 seed weight, in gram), seed length (average of 10 seeds, in cm) were measured for each of the rice lines. The mean of three independent biological replicates was used to calculate the standard error.

### Statistical analysis

Statistical significance was tested by one-way analysis of variance between indicated groups using Student-Newman-Keuls Test using Sigma Plot 12.0 (SPSS, Inc. Chicago, IL, USA). Values with different letters in the bar graph indicate significant differences at P < 0.05 (one-way ANOVA, and the Student–Newman–Keuls tes). Asterisks (*) show the significance at *p < 0.05* and *p < 0.001* mentioned in the figure legend, wherever required.

## RESOURCE AVAILABILITY

The authors declare that all relevant data supporting the findings of the study are available in this article and its Supporting Information files.

## ACKNOWLEDGMENTS

DS and NG acknowledge fellowship from CSIR and DBT, Govt of India, respectively. R.K.C. acknowledges financial support from the DBT-RA programme in Biotechnology and Life Sciences. G.J. acknowledges NIPGR flagship program (102/IFD/SAN/763/2019-20) funded by DBT, Swarna Jayanti Fellowship (SB/SJF/2020-21/01) funded by SERB, Govt of India, and BRIC-NIPGR core research grant. The authors are also thankful to DBT-eLibrary Consortium (DeLCON) for providing access to e-resources.

## AUTHOR CONTRIBUTIONS

GJ conceived the work, supervised and coordinated its progress. DS carried out the detailed characterization of OsNUOR in rice and established its role during the pathogenesis of *R. solani* in rice. RKC performed Y2H and qRT-PCR analysis and both RKC and DS performed GUS reporter as well as BiFC assays. NG assisted in making various constructs. GJ, RKC, and DS have contributed to writing the manuscript and all authors have approved the final manuscript.

## CONFLICT OF INTEREST STATEMENT

No conflict of interest declared.

## FUNDING

The research is supported by NIPGR flagship program (102/IFD/SAN/763/2019-20) funded by DBT, Govt of India, Swarna Jayanti Fellowship (SB/SJF/2020-21/01) funded by SERB, Govt of India, and BRIC-NIPGR core research grant.

## Supplementary Files

**Figure S1:**
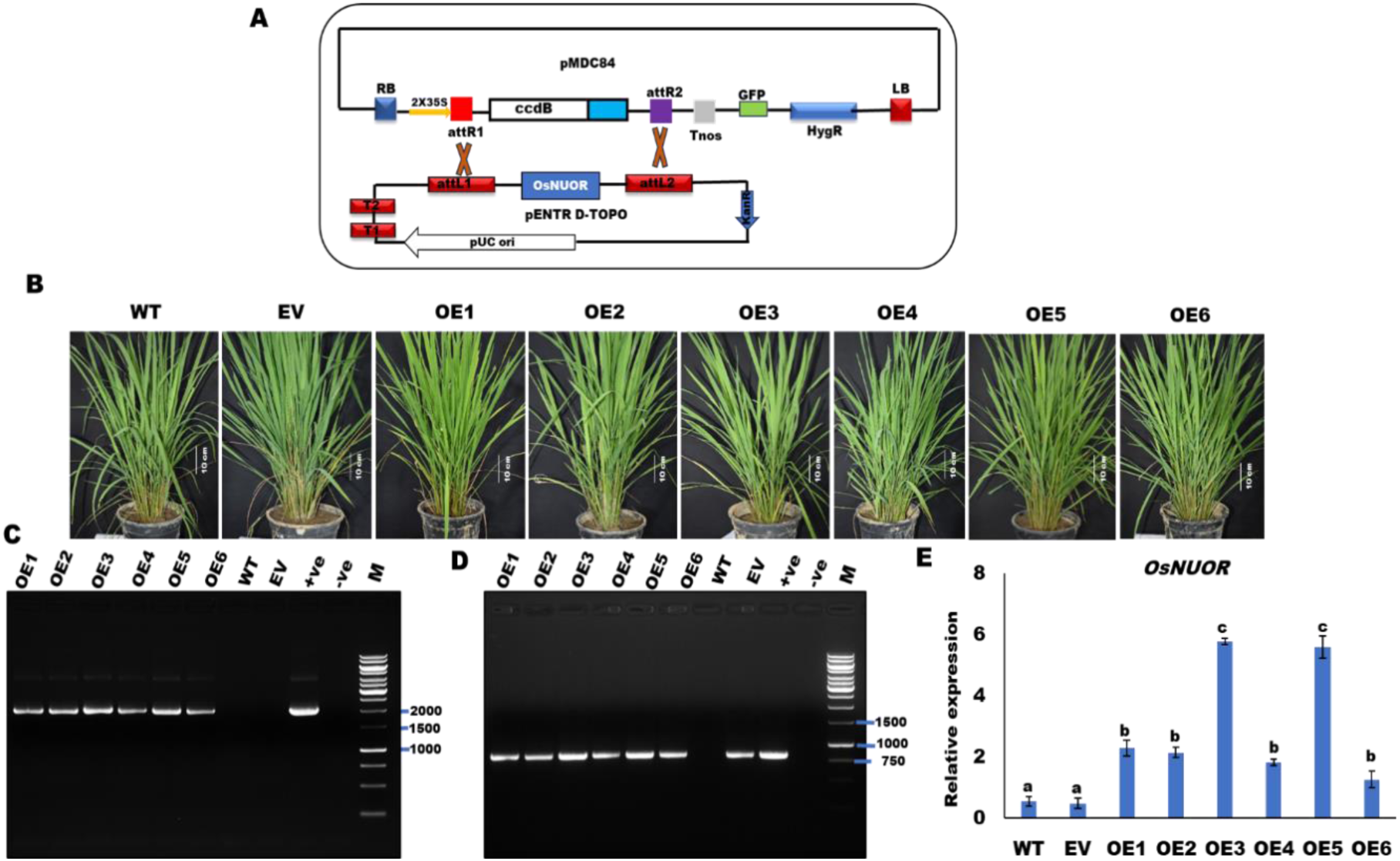
Development of *OsNUOR* overexpression rice lines. **(A)** T-DNA map of pMDC84 gateway binary vector used to develop *OsNUOR* overexpression (OE) and empty vector (EV) lines. **(B)** Growth phenotype of 45-day-grown OE and EV lines. **(C)** The presence of the transgene in T_2_ lines was confirmed by PCR using 35S-forward and gene reverse primer. The amplicon size of 2152 bp confirms the presence of transgene. **(D)** The presence of T-DNA was confirmed by PCR amplification of the hygromycin B phosphotransferase (*HPT*) gene. The amplicon size of 809 bp suggests the presence of T-DNA. **(E)** The relative expression of *OsNUOR* in different rice lines, quantified using rice *actin* gene as an endogenous control. The bar denotes the average of three independent biological replicates and the error bar represents mean ± standard error. Different letters represent a significant difference between samples at *p* < 0.05 (one-way ANOVA, Student-Newman-Keuls test).

**Figure S2:**
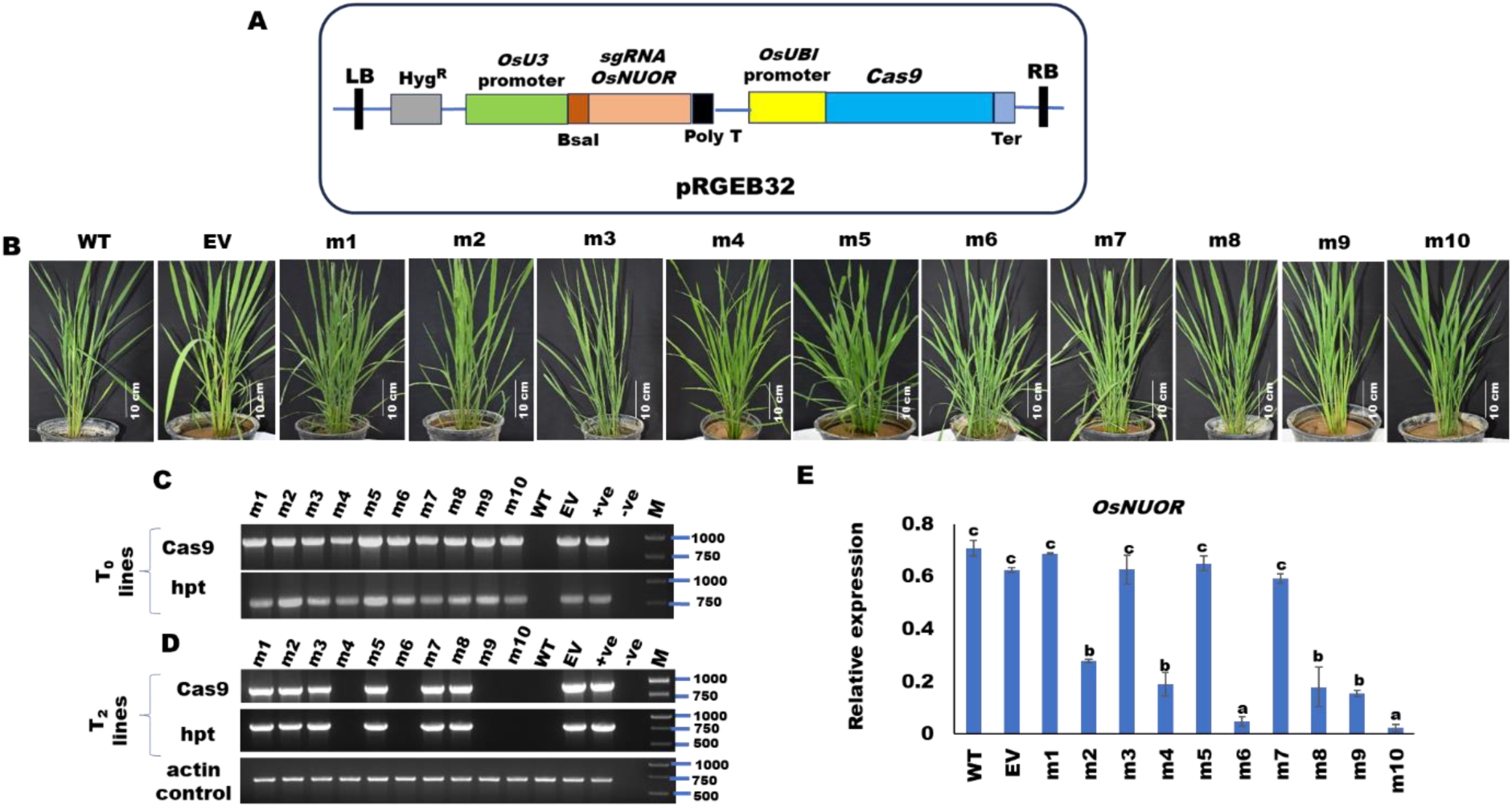
Development of *OsNUOR*-edited rice lines. **(A)** Map of pRGEB32 binary vector that was used for guide RNA constructs for genome editing of *OsNUOR*. **(B)** The representative photographs show the growth phenotype of OsNUOR-edited (m1-m10), wild-type (WT), and empty vector (EV) lines. **(C)** The presence of T-DNA in T_0_ lines. The PCR amplicon of 959 bp reflects the presence of Cas9, while 809 bp amplicon reflects the presence of hygromycin B phosphotransferase (hpt) gene. **(D)** The presence of T-DNA (Cas9 and hpt) in T_2_ lines. Rice *actin* gene (amplicon size of 698 bp) was used as a DNA control. **(E)** Graph shows the relative expression of *OsNUOR* in different rice lines. Relative expression was quantified using rice *actin* gene as an endogenous control. The bar denotes the average of three independent biological replicates and the error bar represents mean ± standard error. Different letters represent a significant difference between samples at *p* < 0.05 (one-way ANOVA, Student-Newman-Keuls test).

**Figure S3:**
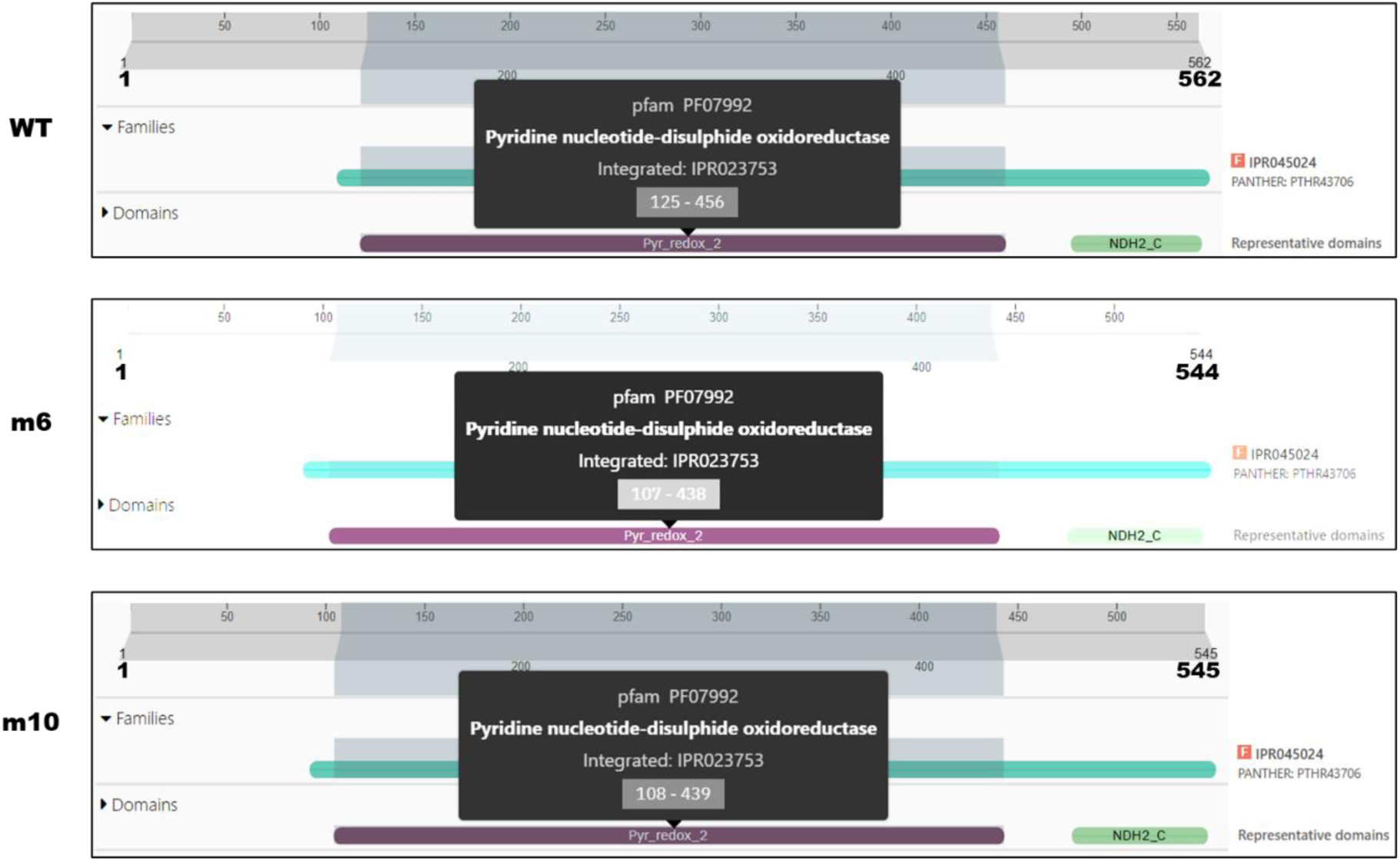
The functional domain of OsNUOR remained intact in the edited line. Schematics of Interpro scan output, reflecting that both wild-type and edited lines (m6 and m10) have intact functional domain i.e pyridine nucleotide-disulfide oxidoreductase [Pyr_redox2 domain (pfam PF07992)] belonging to NADH dehydrogenase family (PANTHER ID: PTHR43706).

**Figure S4:**
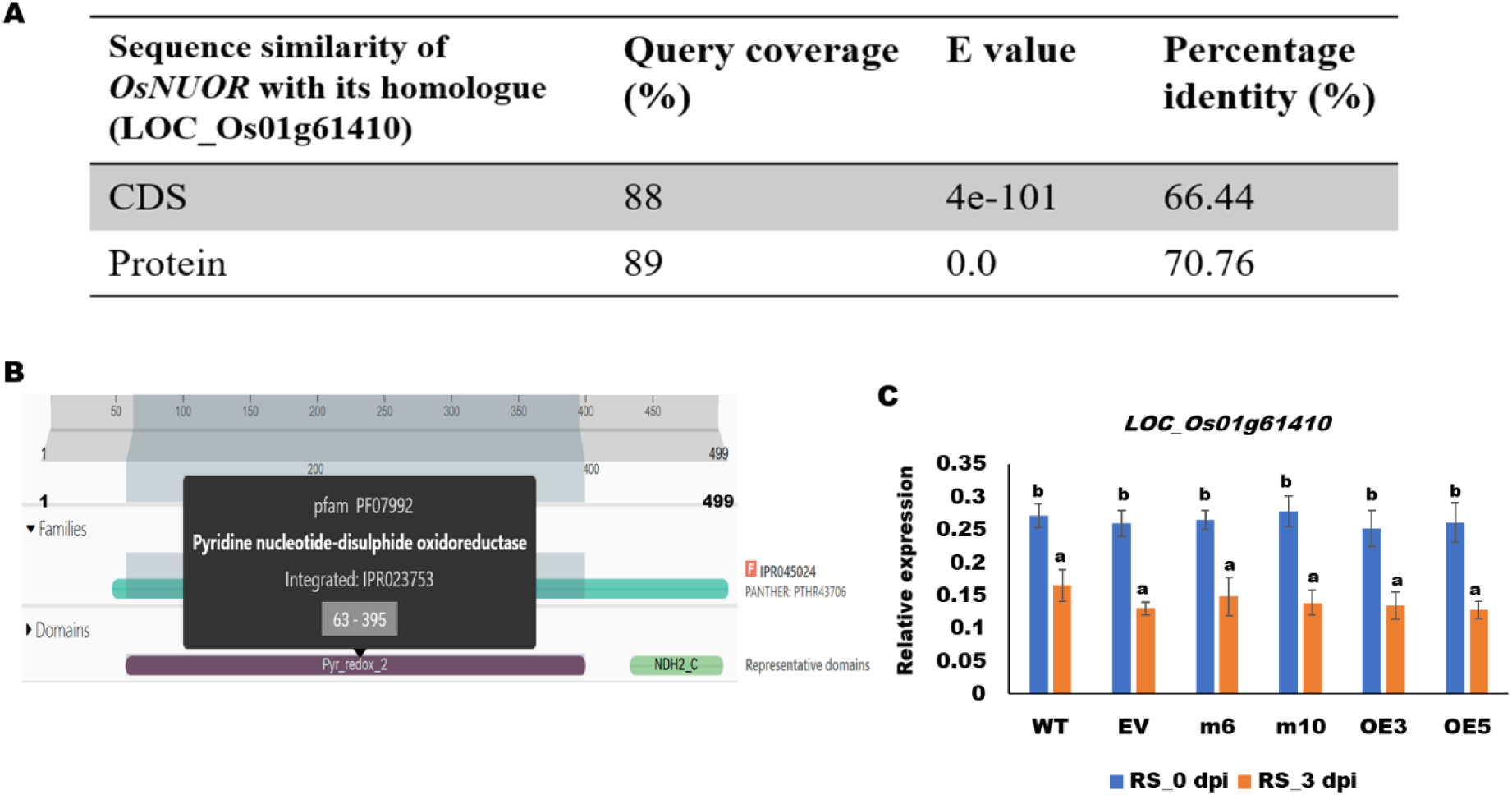
Off-target editing is not observed in OsNUOR homologue gene in rice. **(A)** Sequence similarity of *OsNUOR* with its homologue LOC_Os01g61410 in rice. **(B)** Schematics of Interpro scan output, reflecting that LOC_Os01g61410 harbours pyridine nucleotide-disulfide oxidoreductase [Pyr_redox2 domain (pfam PF07992)] domain belonging to NADH dehydrogenase family (PANTHER ID: PTHR43706). **(C)** Relative expression of *LOC_Os01g61410* is comparable in different rice lines. Relative expression was quantified using rice *actin* gene as an endogenous control. The bar denotes the average of three independent biological replicates and the error bar represents mean ± standard error. Different letters represent a significant difference between samples at *p* < 0.0*5* (one-way ANOVA, Student-Newman-Keuls test).

**Figure S5:**
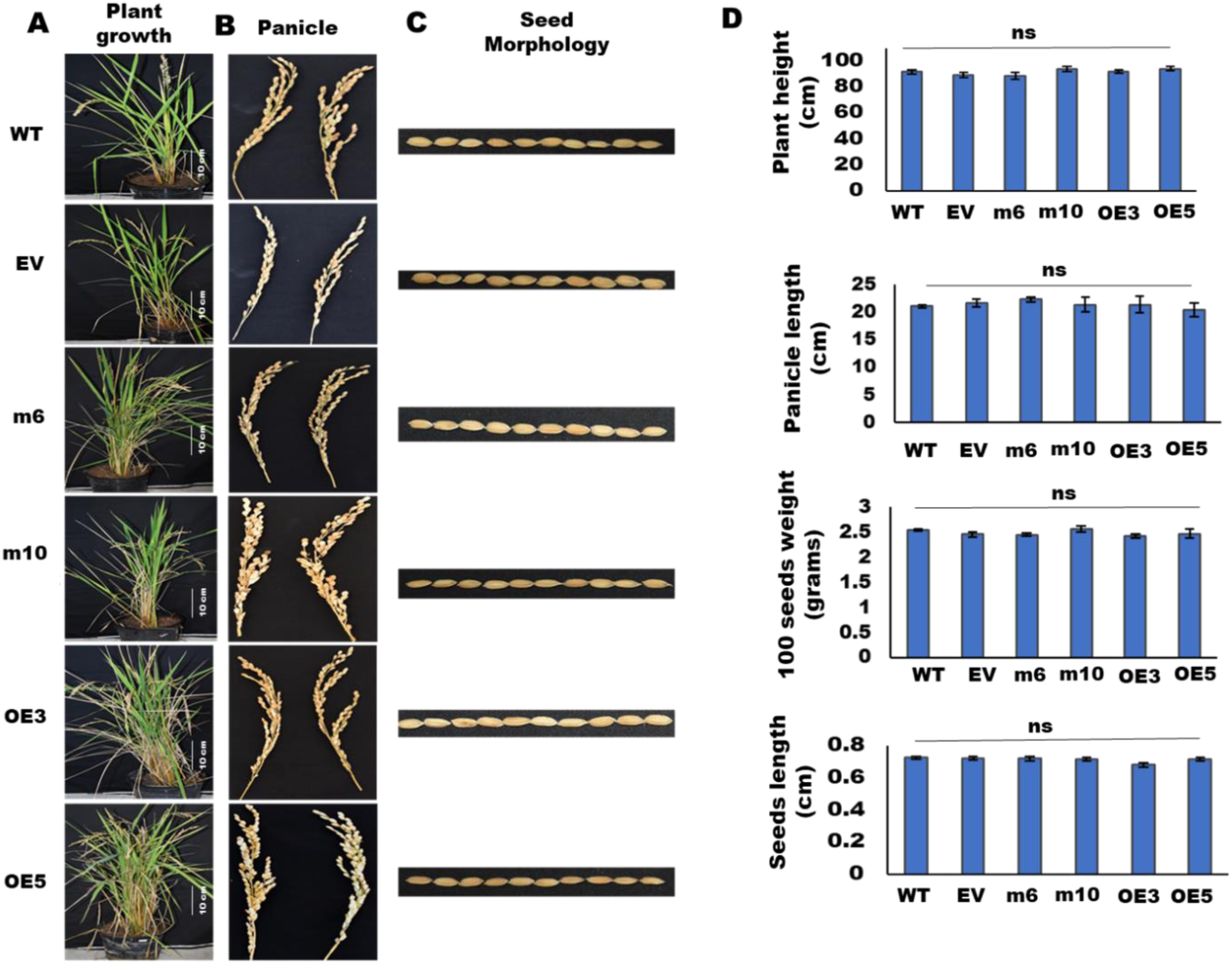
Agronomical traits are comparable in different rice lines. **(A)** Plant growth **(B)** Panicle and **(C)** Seed morphology traits of different rice lines. The quantitative measurement of various agronomical traits, **(D)** Plant height (n=20), panicle length (n=20), 100 seed weight (n=10), and seed length (n=10) in different rice lines. The bar denotes the average of three independent biological replicates and the error bar represents mean ± standard error. ns indicates no significant difference.

**Figure S6:**
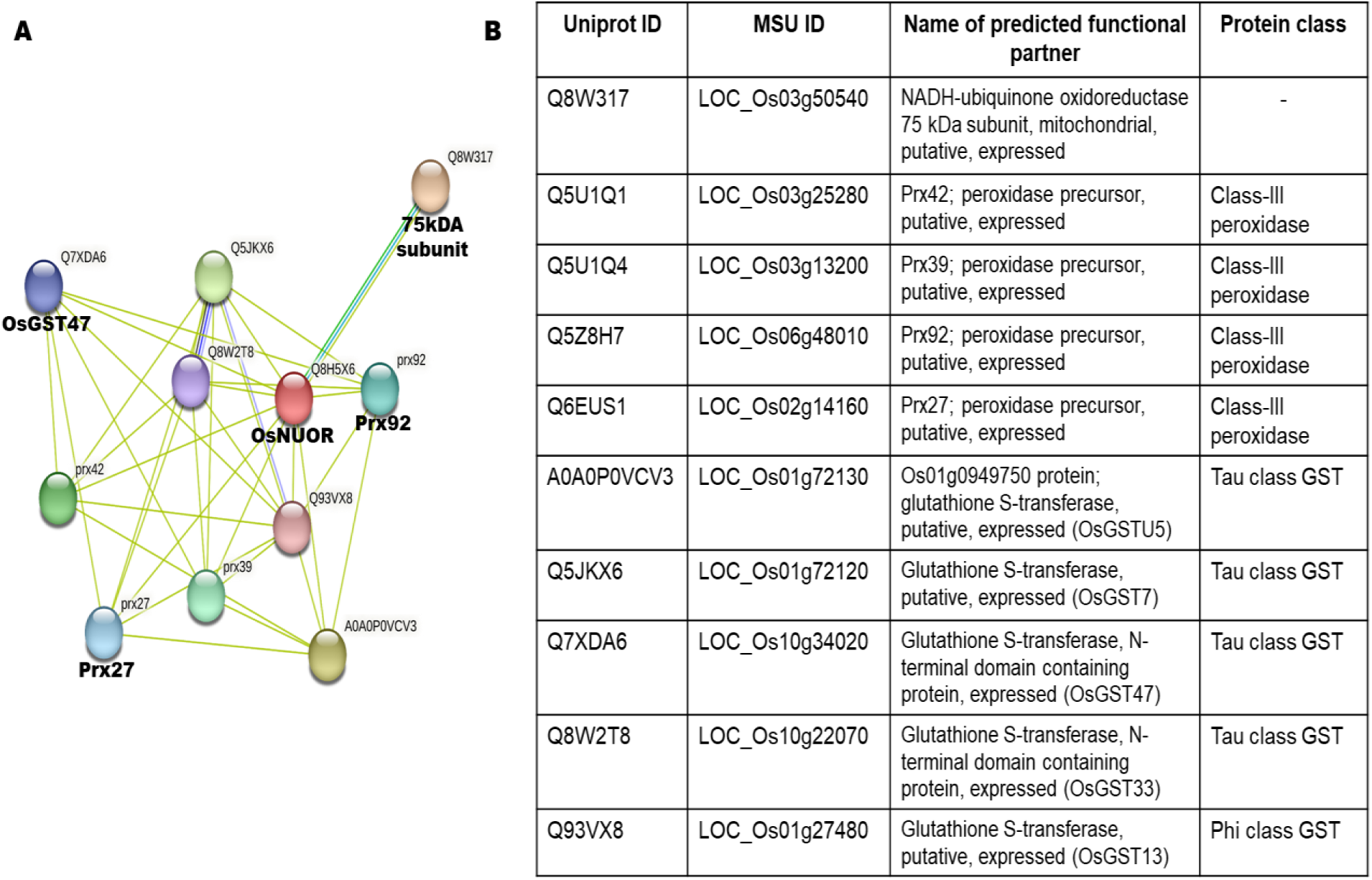
STRING analysis reflecting potential interacting partners of OsNUOR. **(A)** Plot showing the interaction of OsNUOR (Q8H5X6) with partners in STRING database version 11.5. **(B)** The predicted interacting partners are tabulated.

**Figure S7:**
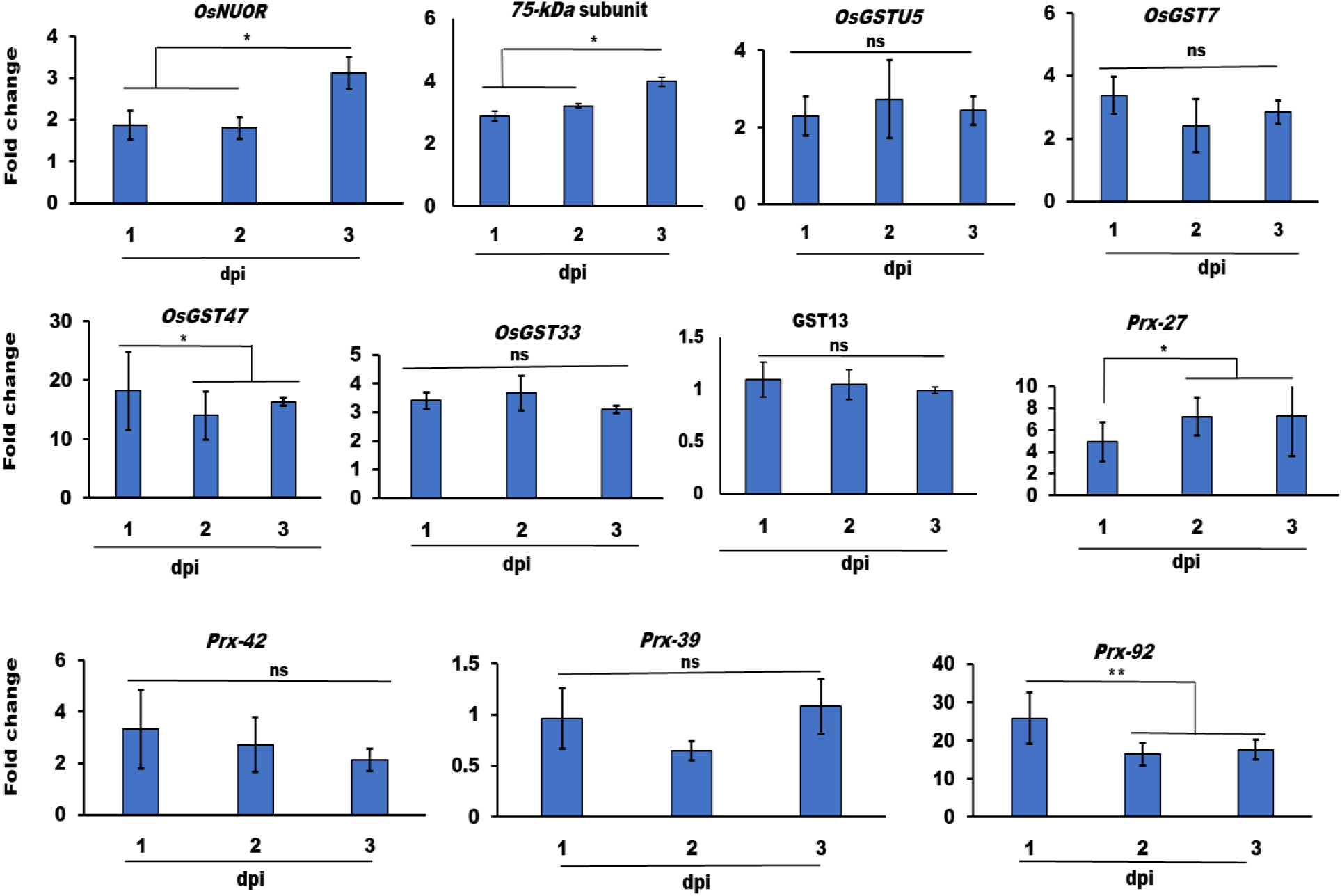
The interacting partners of OsNUOR are upregulated under *R. solani*-infected conditions in rice. qRT-PCR-based estimation of fold change in gene expression, with respect to 0 dpi samples, using rice *actin* gene as endogenous control. The bar denotes the average of three independent biological replicates and the error bar represents mean ± standard error. ‘*’ indicate significant differences at *p* < 0.05 (estimated using one-way ANOVA and Student-Newman-Keuls test). ns indicates no significant difference.

**Figure S8:**
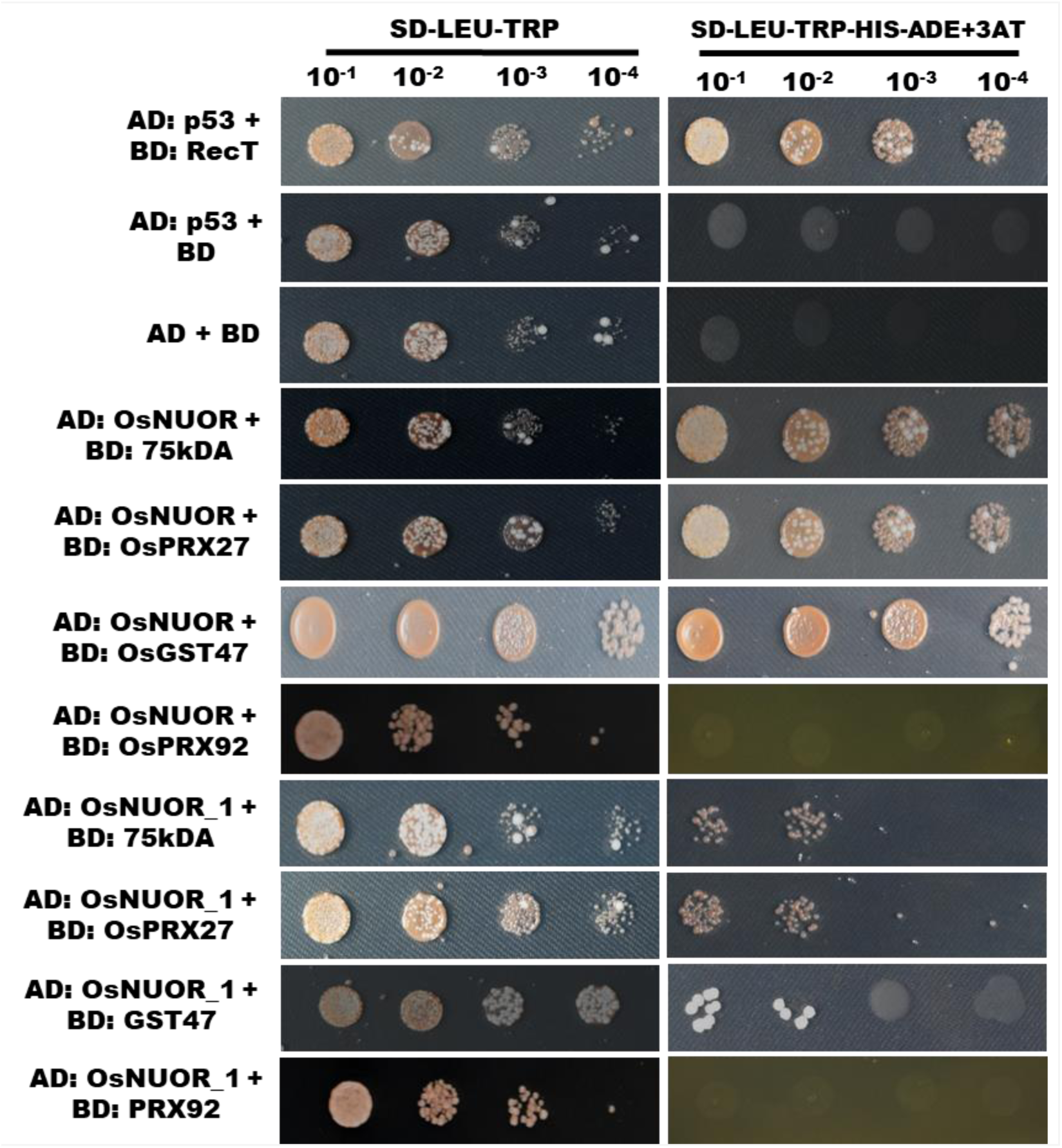
Yeast two-hybrid assay to test the interaction of OsNUOR with predicted target proteins. Y2H assay reflecting the interaction of OsNUOR/OsNUOR_1 with target proteins (75 kDa subunit of complex-1, Prx27, and GST47). The growth of yeast cells on quadruple drop-out plates (SD-Leu-Trp-Ade-His), reflects positive interactions. AD is prey vector (pGADT7) and BD is bait vector (pGBKT7). 3-AT; 3-aminotrizole (15 mM). AD: p53 + BD: RecT is used as positive control and AD: p53 +BD is used as negative control.

**Figure S9:**
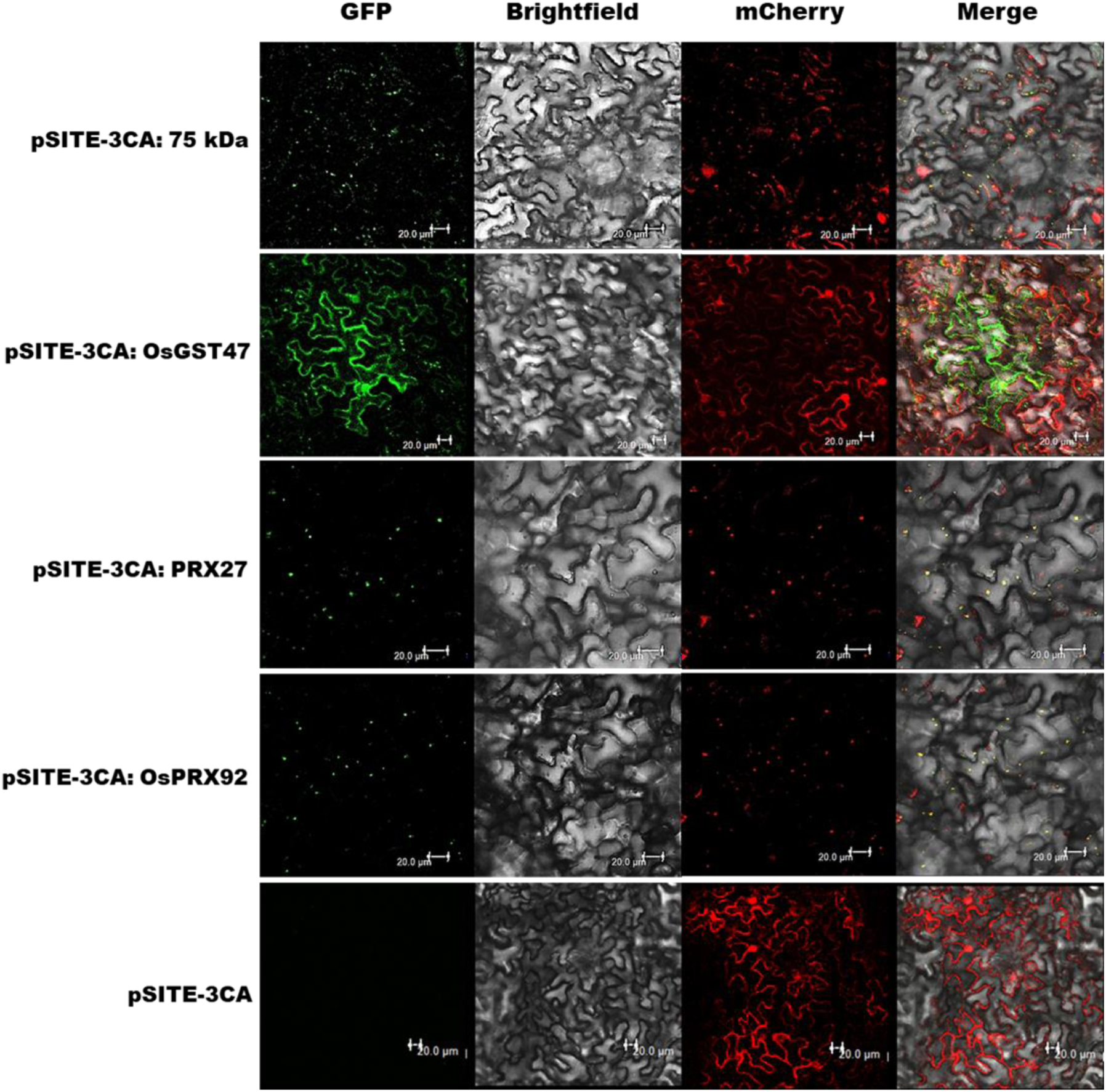
Localization of selected interacting partners of OsNUOR in *N. benthamiana*. Confocal images showing localization of YFP-tagged OsGST47 in the cytosol and YFP-tagged OsPRX27, OsPRX92 and Os75 kDa subunit of complex-1 in mitochondria (being colocalized with mCherry-tagged mitochondrial marker, mt-rk CD3-991). The YFP was visualized under GFP (green fluorescent protein) filter, while mCherry was visualized under RFP (red fluorescent protein) filter. Scale bars = 20 μm.

**Figure S10:**
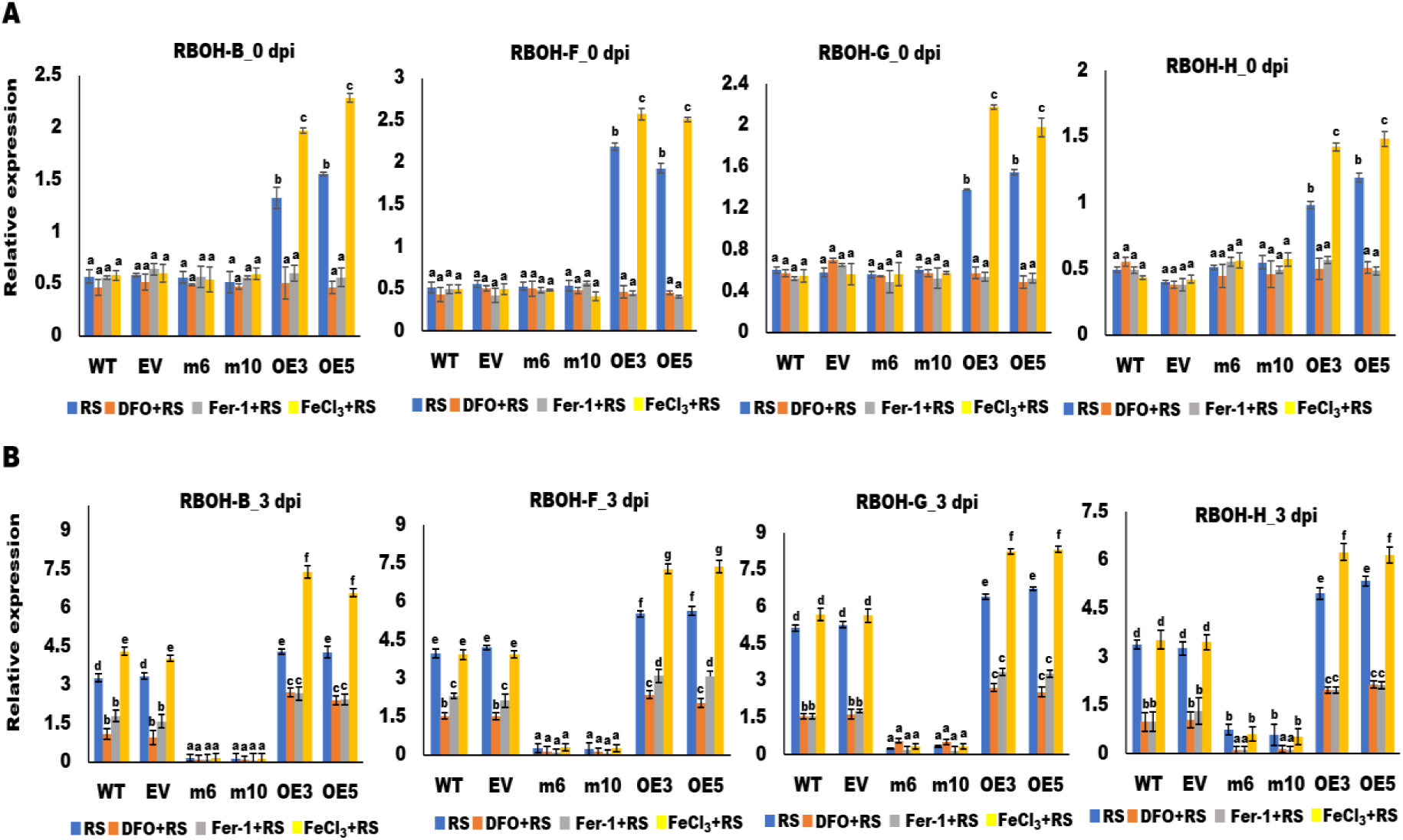
Editing of *OsNUOR* compromises the expression of ROS-responsive genes. Relative expression of *RBOH* homologs (*OsRBOH-B*, *OsRBOH-F*, *OsRBOH-G* and *OsRBOH-H*) in different rice lines with and without DFO/FER-1/FeCl_3_ treatment at **(A)** 0 dpi and **(B)** 3 dpi of *R. solani* infection. Relative expression was quantified using *actin* gene of *Oryza sativa* as an endogenous control. The bar denotes the average of three independent biological replicates and the error bar represents mean ± standard error. Different letters represent a significant difference between samples at *p* < 0.05 (one-way ANOVA, Student-Newman-Keuls test).

**Figure S11:**
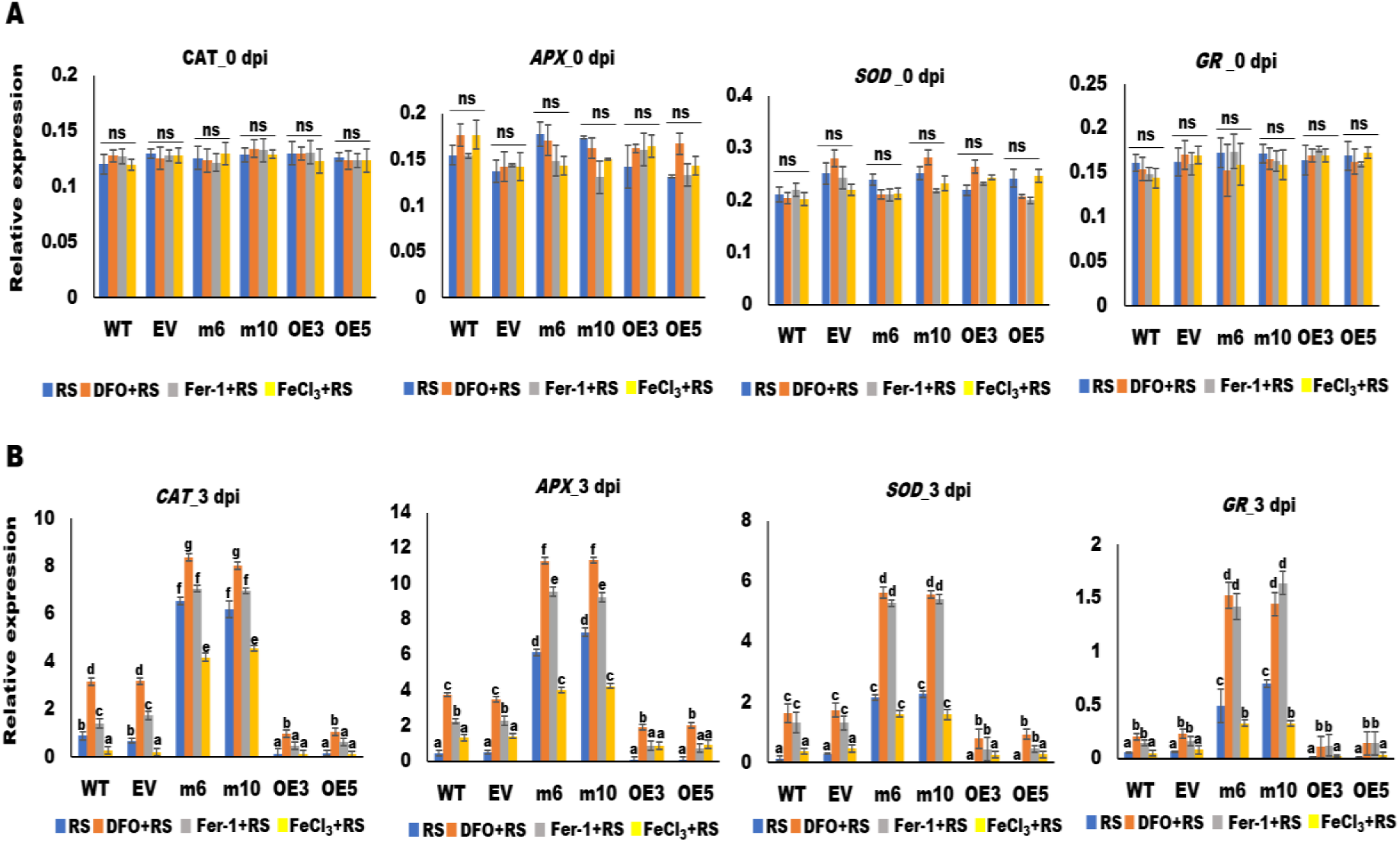
Antioxidant defense is induced in edited lines, under *R. solani*-infected conditions. Relative expression of antioxidant genes (*OsCAT*, *OsAPX*, *OsSOD* and *OsGR*) in different rice lines, with and without DFO/FER-1/FeCl_3_ treatment at **(A)** 0 dpi and **(B)** 3 dpi of *R. solani* infection. Relative expression was quantified using rice *actin* gene as an endogenous control. The bar denotes the average of three independent biological replicates and the error bar represents mean ± standard error. Different letters represent a significant difference between samples at *p* < 0.05 (one-way ANOVA, Student-Newman-Keuls test). ns indicates no significant difference.

**Figure S12:**
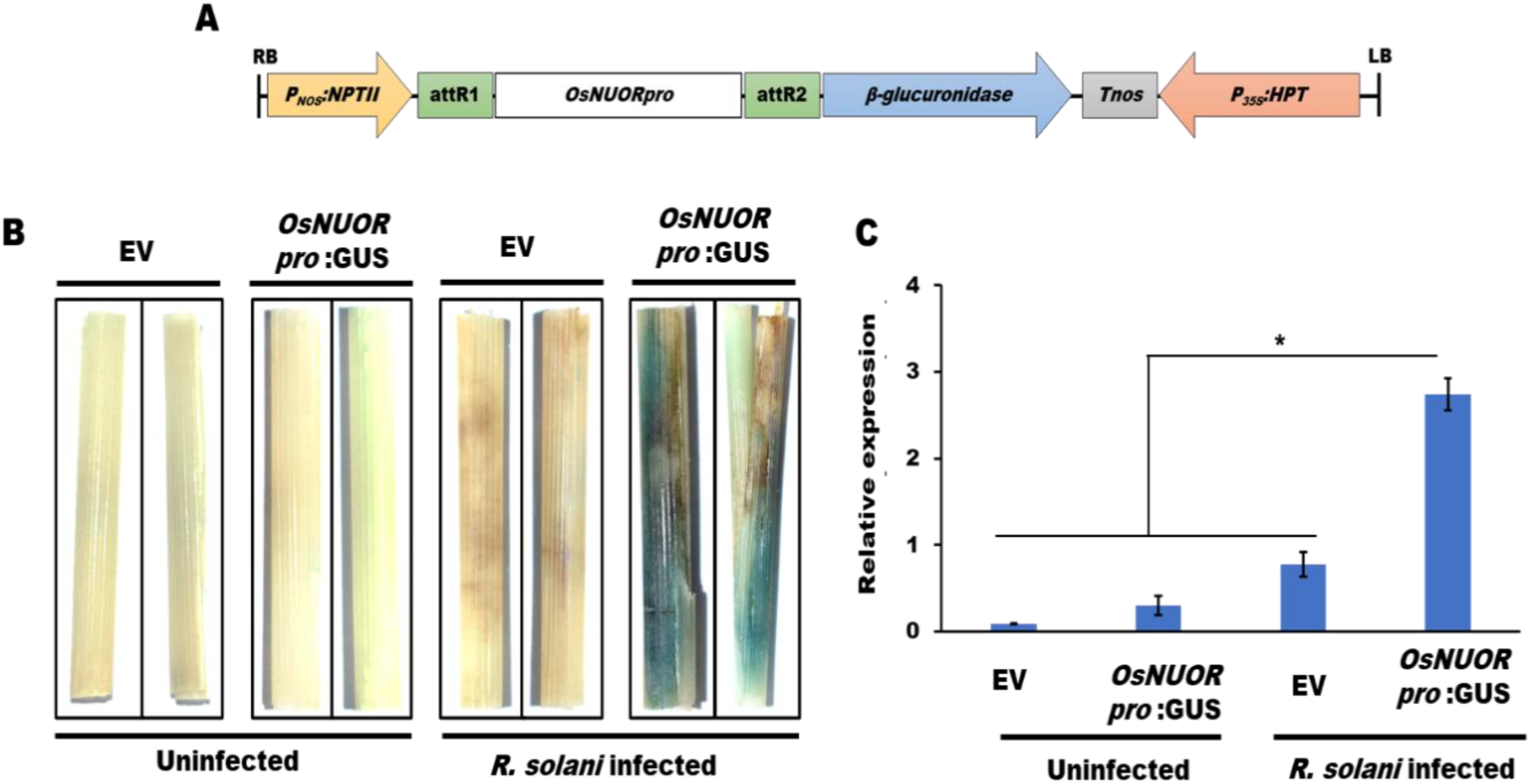
*OsNUOR* is upregulated upon *R. solani* infection in rice. **(A)** The map of pGWB3 gateway vector used for cloning of *OsNUOR* promoter for reporter GUS assay. **(B)** β-glucuronidase (GUS) assay reflecting activation of *OsNUOR* promoter (appearance of blue colour) in rice sheaths upon *R. solani* infection at 3 dpi. **(C)** qRT-PCR-based quantification of β-glucuronidase (*GUS*) gene expression using rice *actin* gene as an endogenous control. The bar denotes the average of three independent biological replicates and the error bar represents mean ± standard error. ‘*’ indicate significant differences at *p* < 0.05 (estimated using one-way ANOVA and Student-Newman-Keuls test).

**Supplementary table 1:**
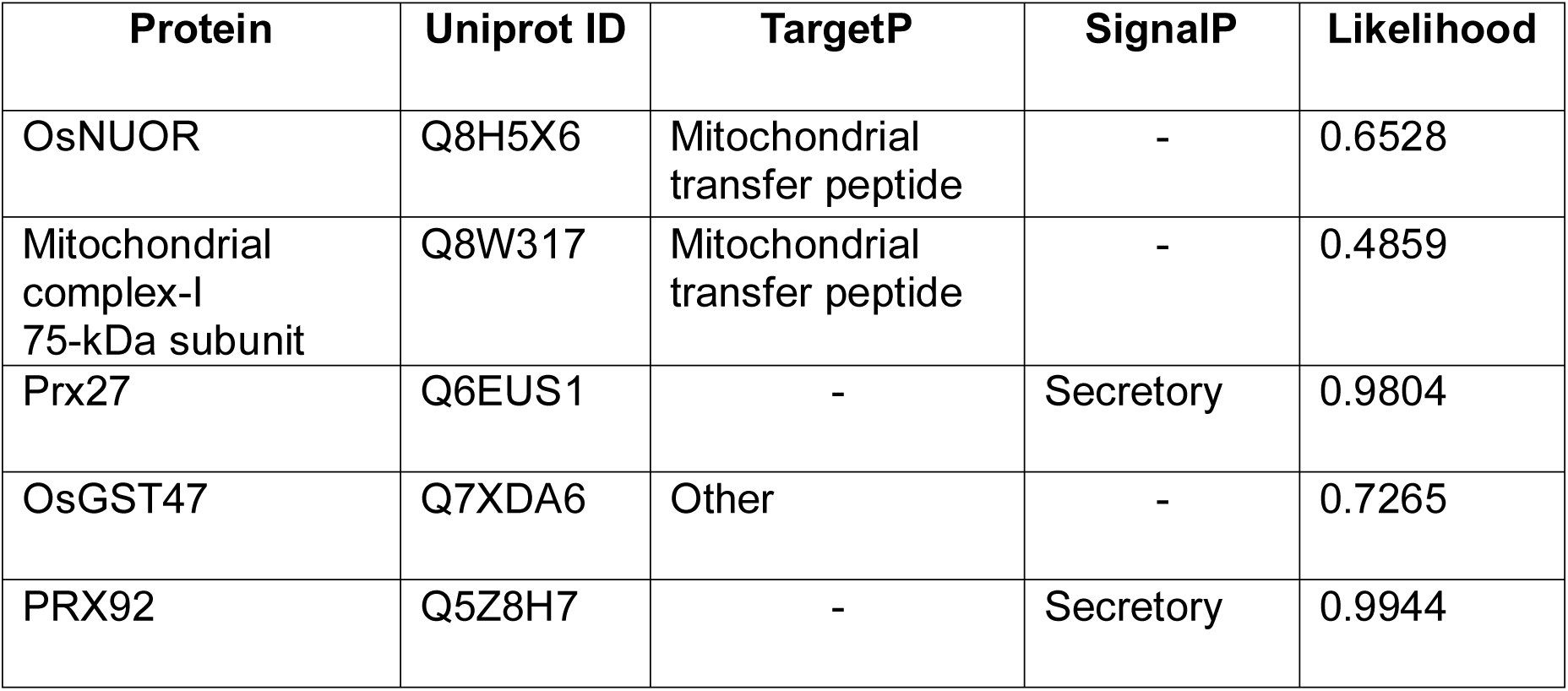
TargetP and SignalP results of *OsNUOR* and its interacting partners.

**Supplementary table 2:**
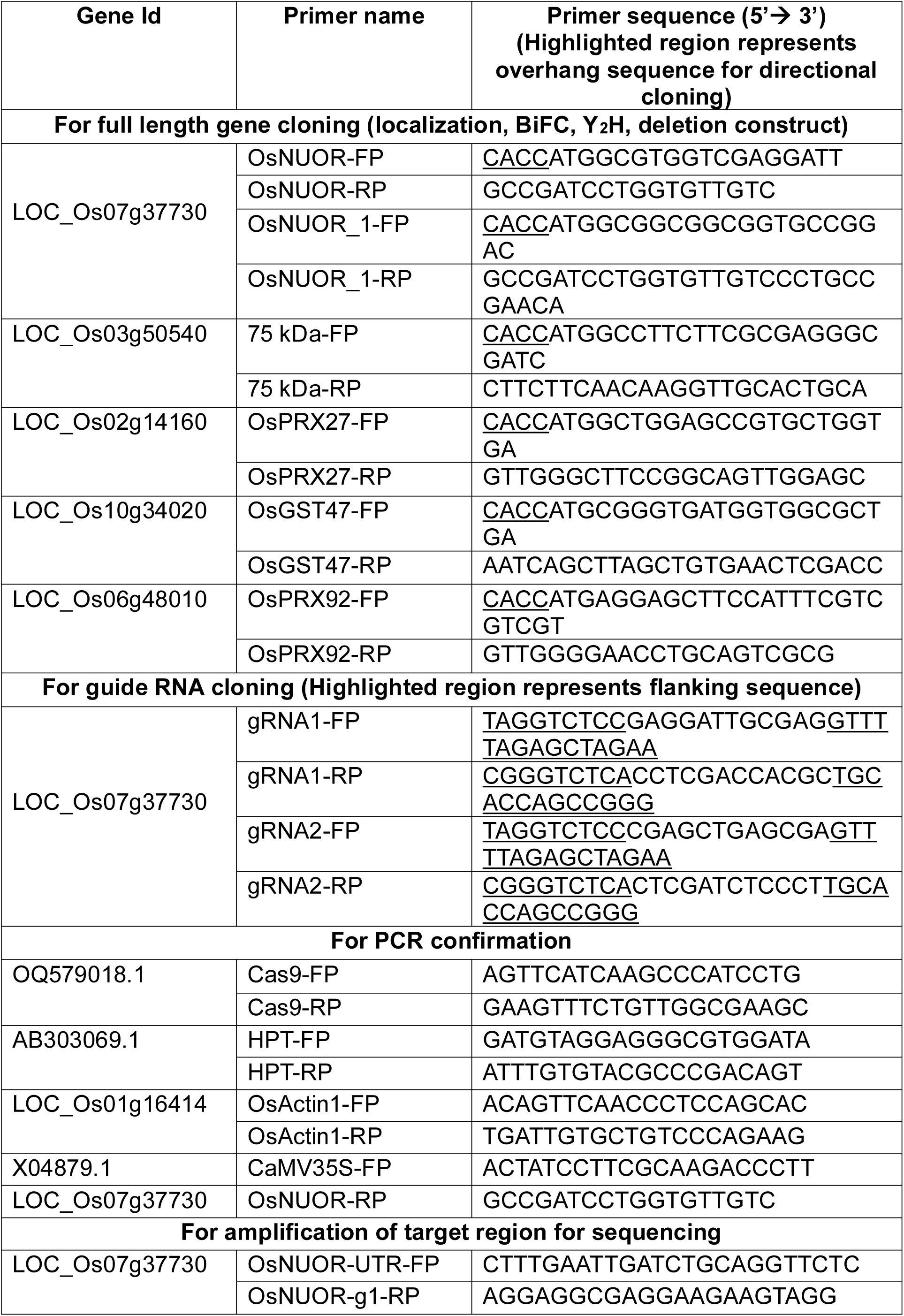
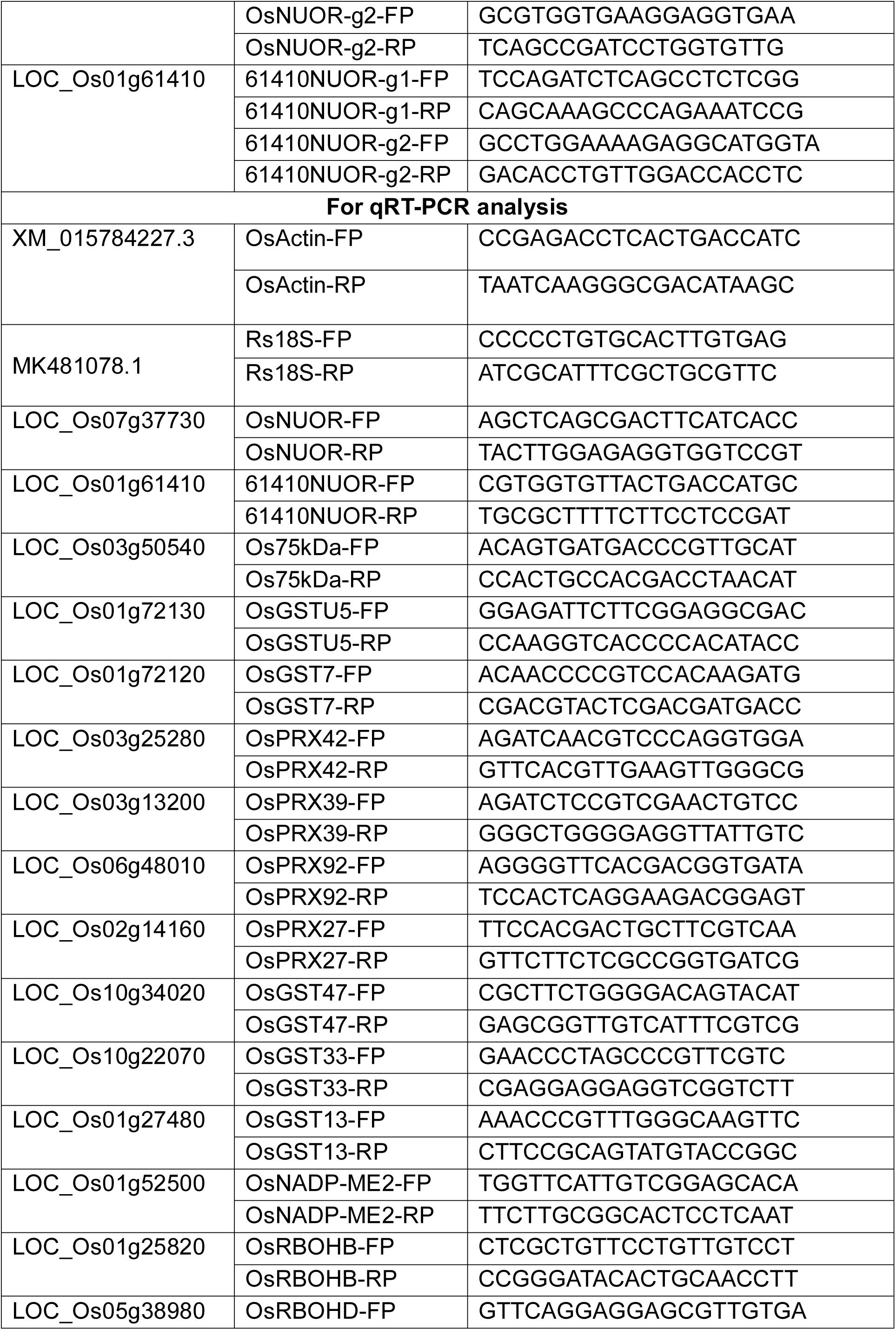
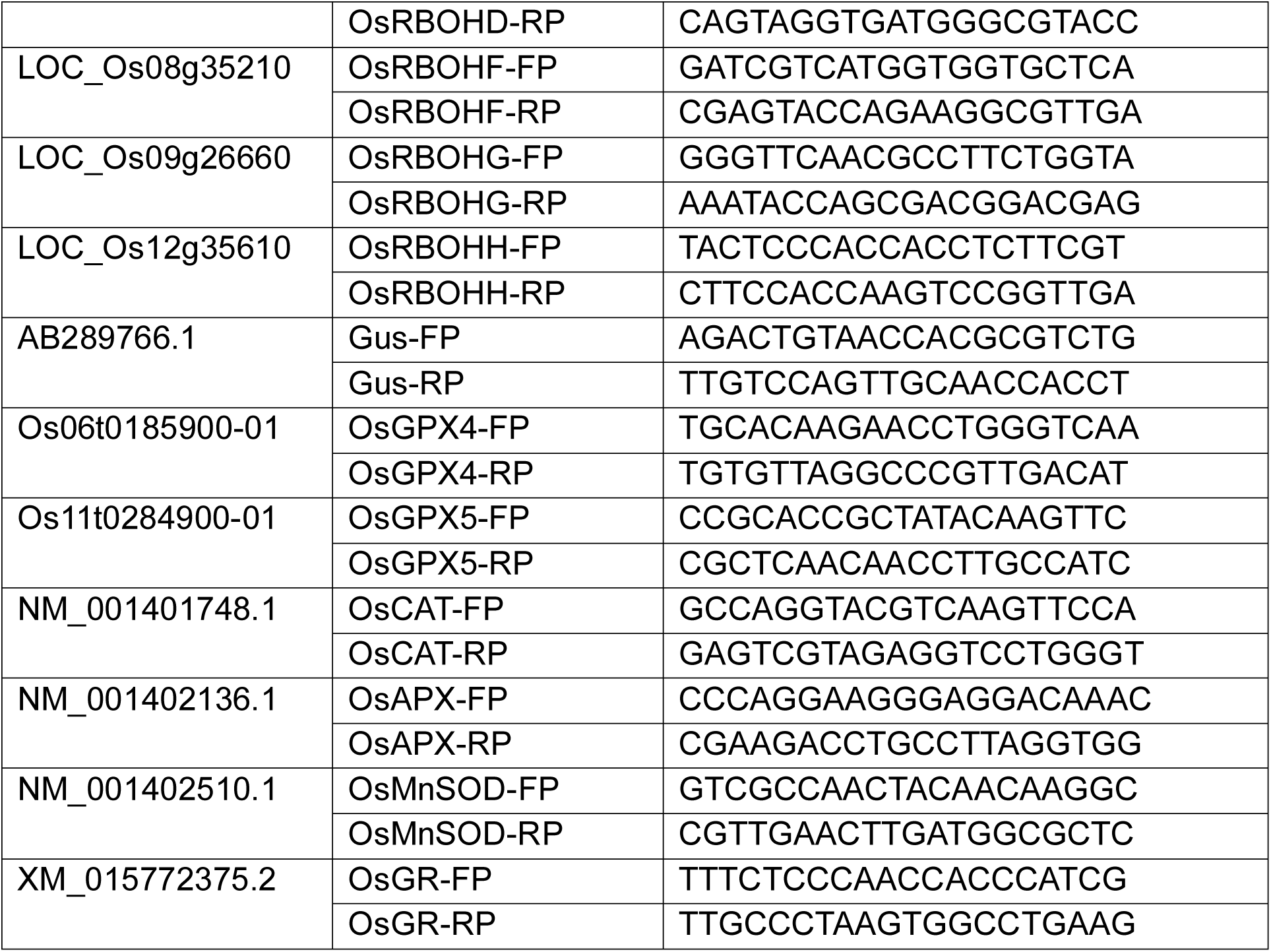
List of primers used in the study.

## Notes

### Competing Interest Statement

The authors have declared no competing interest.

