## Supplementary Files for "Alternative NADH:ubiquinone oxidoreductase modulates disease susceptibility in rice by interfering ROS homeostasis and ferroptosis"

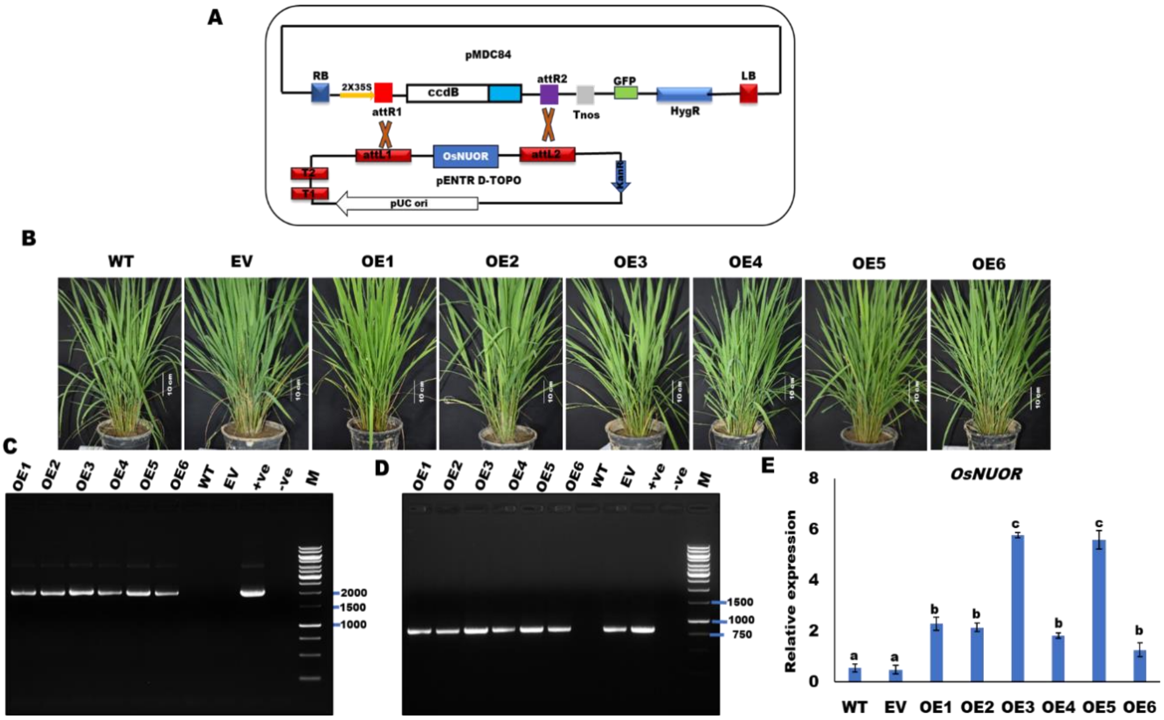


**Figure S1: Development of *OsNUOR* overexpression rice lines (A)** T-DNA map of pMDC84 gateway binary vector used to develop *OsNUOR* overexpression (OE) and empty vector (EV) lines. **(B)** Growth phenotype of 45-day-grown OE and EV lines. **(C)** The presence of the transgene in $T_2$ lines was confirmed by PCR using 35S-forward and gene reverse primer. The amplicon size of 2152 bp confirms the presence of transgene. **(D)** The presence of T-DNA was confirmed by PCR amplification of the hygromycin B phosphotransferase (*HPT*) gene. The amplicon size of 809 bp suggests the presence of T-DNA. **(E)** The relative expression of *OsNUOR* in different rice lines, quantified using rice *actin* gene as an endogenous control. The bar denotes the average of three independent biological replicates and the error bar represents mean ± standard error. Different letters represent a significant difference between samples at $p < 0.05$ (one-way ANOVA, Student-Newman-Keuls test).

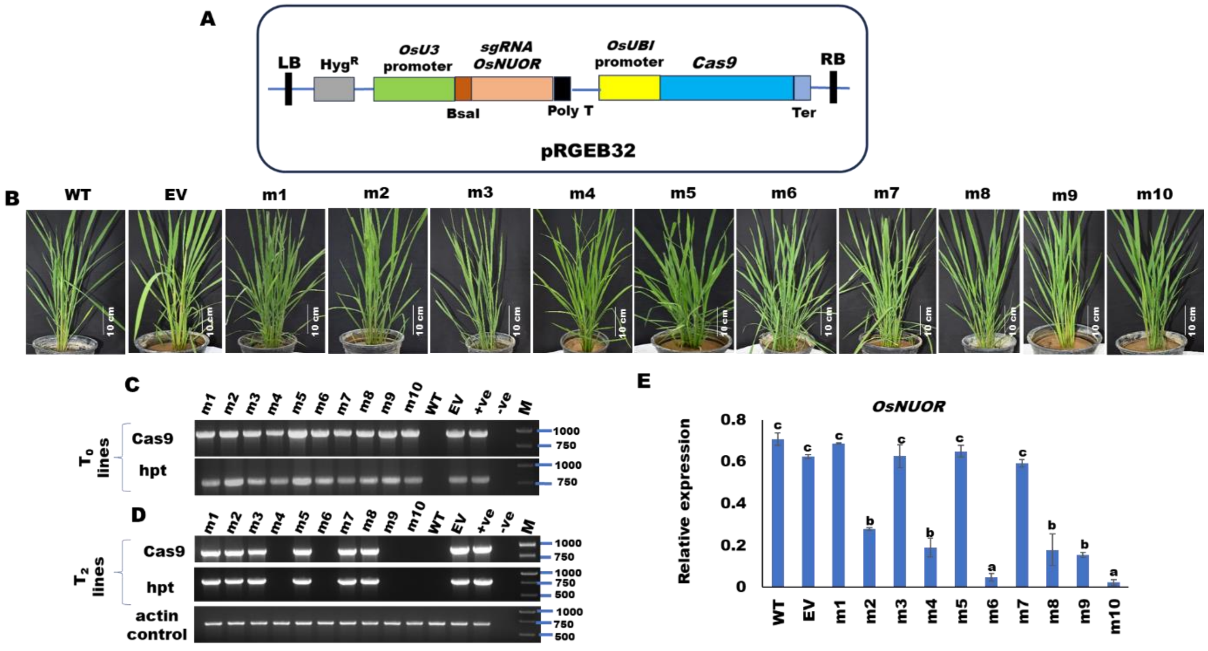


**Figure S2: Development of *OsNUOR*-edited rice lines. (A)** Map of pRGEB32 binary vector that was used for guide RNA constructs for genome editing of *OsNUOR*. **(B)** The representative photographs show the growth phenotype of OsNUOR-edited (m1-m10), wild-type (WT), and empty vector (EV) lines. **(C)** The presence of T-DNA in $T_0$ lines. The PCR amplicon of 959 bp reflects the presence of Cas9, while 809 bp amplicon reflects the presence of hygromycin B phosphotransferase (hpt) gene. **(D)** The presence of T-DNA (Cas9 and hpt) in $T_2$ lines. Rice *actin* gene (amplicon size of 698 bp) was used as a DNA control. **(E)** Graph shows the relative expression of *OsNUOR* in different rice lines. Relative expression was quantified using rice *actin* gene as an endogenous control. The bar denotes the average of three independent biological replicates and the error bar represents mean ± standard error. Different letters represent a significant difference between samples at $p < 0.05$ (one-way ANOVA, Student-Newman-Keuls test).

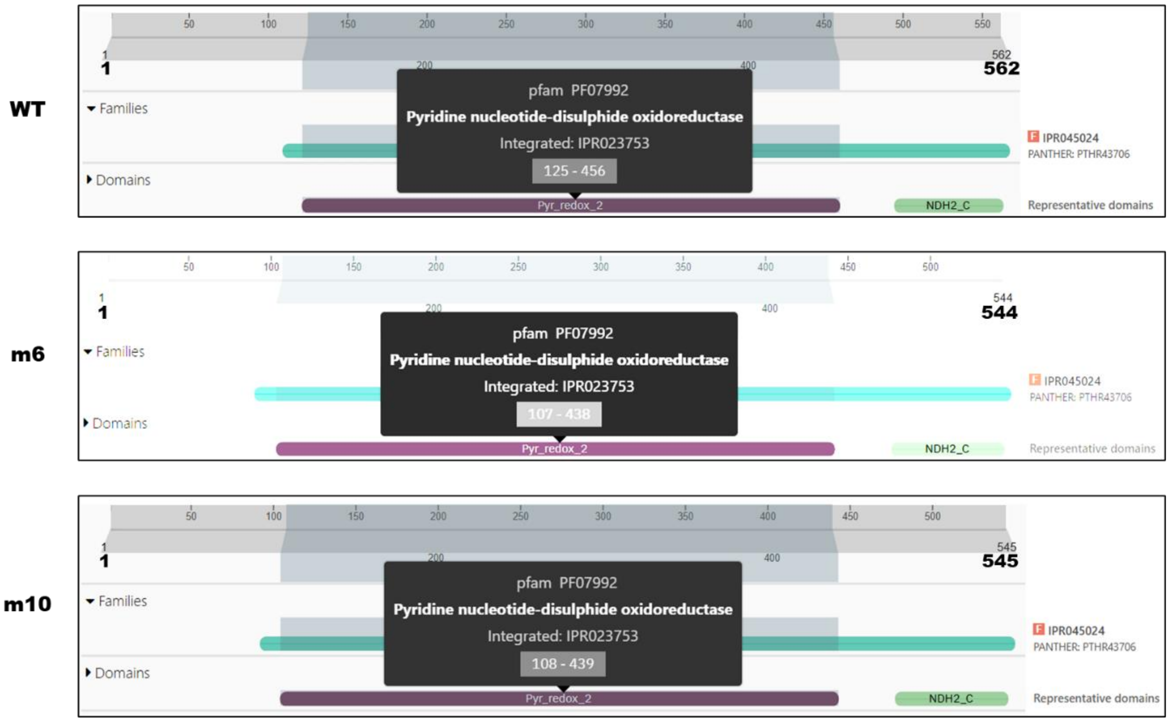


**Figure S3: The functional domain of OsNUOR remained intact in the edited line.** Schematics of Interpro scan output, reflecting that both wild-type and edited lines (m6 and m10) have intact functional domain i.e pyridine nucleotide-disulfide oxidoreductase [Pyr_redox2 domain (pfam PF07992)] belonging to NADH dehydrogenase family (PANTHER ID: PTHR43706).

A

| Sequence similarity of *OsNUOR* with its homologue (LOC_Os01g61410) | Query coverage (%) | E value | Percentage identity (%) |
|---|---|---|---|
| CDS | 88 | 4e-101 | 66.44 |
| Protein | 89 | 0.0 | 70.76 |

B C

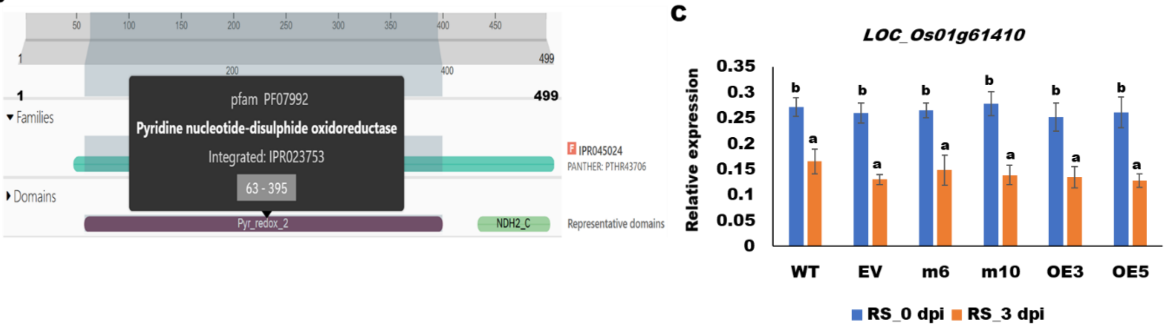


**Figure S4: Off-target editing is not observed in OsNUOR homologue gene in rice. (A)** Sequence similarity of *OsNUOR* with its homologue LOC_Os01g61410 in rice. **(B)** Schematics of Interpro scan output, reflecting that LOC_Os01g61410 harbours pyridine nucleotide-disulfide oxidoreductase [Pyr_redox2 domain (pfam PF07992)] domain belonging to NADH dehydrogenase family (PANTHER ID: PTHR43706). **(C)** Relative expression of *LOC_Os01g61410* is comparable in different rice lines. Relative expression was quantified using rice *actin* gene as an endogenous control. The bar denotes the average of three independent biological replicates and the error bar represents mean ± standard error. Different letters represent a significant difference between samples at $p$ < 0.05 (one-way ANOVA, Student-Newman-Keuls test).

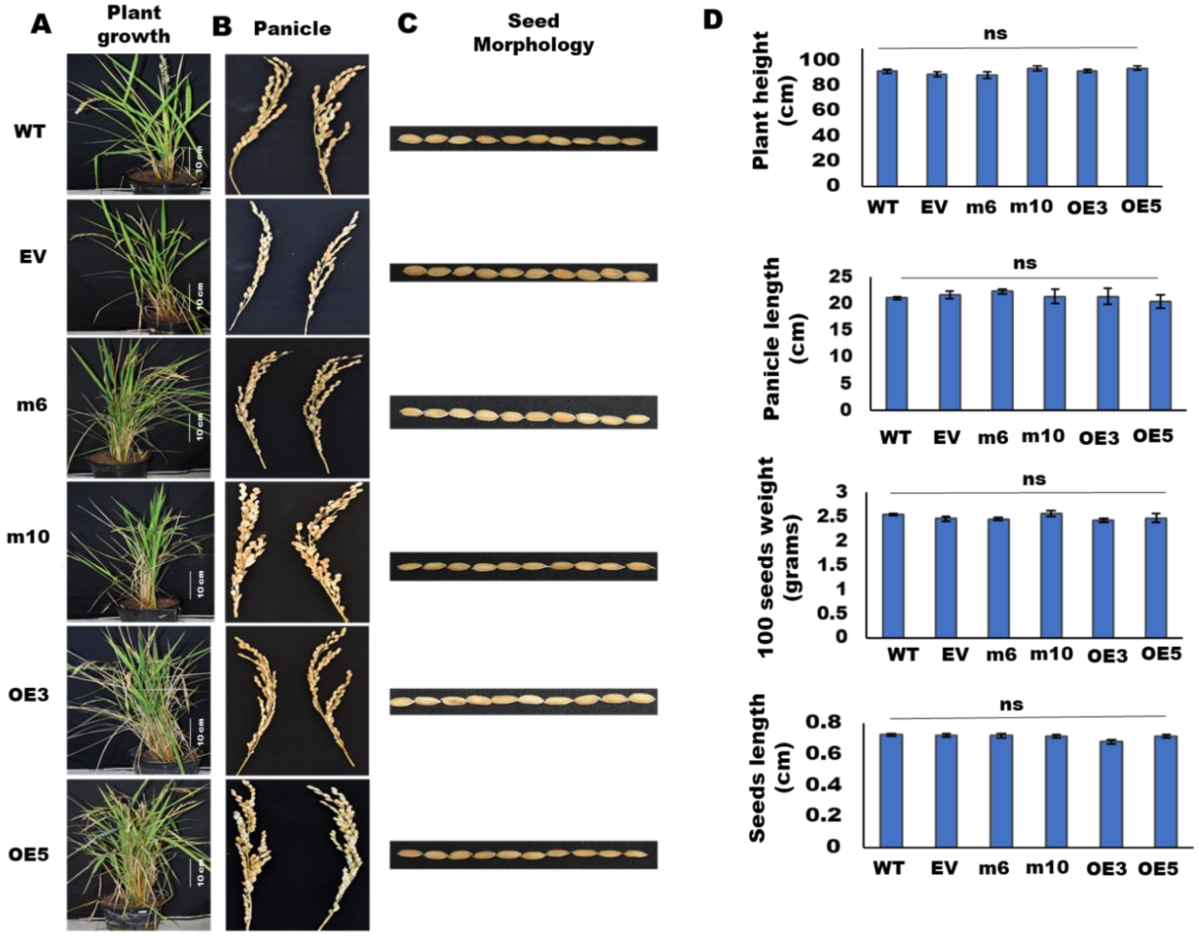


**Figure S5: Agronomical traits are comparable in different rice lines. (A)** Plant growth **(B)** Panicle and **(C)** Seed morphology traits of different rice lines. The quantitative measurement of various agronomical traits, **(D)** Plant height (n=20), panicle length (n=20), 100 seed weight (n=10), and seed length (n=10) in different rice lines. The bar denotes the average of three independent biological replicates and the error bar represents mean ± standard error. ns indicates no significant difference.

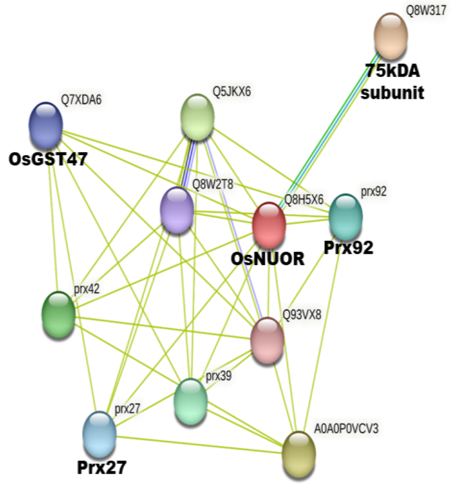


B

| Uniprot ID | MSU ID | Name of predicted functional partner | Protein class |
|---|---|---|---|
| Q8W317 | LOC_Os03g50540 | NADH-ubiquinone oxidoreductase 75 kDa subunit, mitochondrial, putative, expressed | - |
| Q5U1Q1 | LOC_Os03g25280 | Prx42; peroxidase precursor, putative, expressed | Class-III peroxidase |
| Q5U1Q4 | LOC_Os03g13200 | Prx39; peroxidase precursor, putative, expressed | Class-III peroxidase |
| Q5Z8H7 | LOC_Os06g48010 | Prx92; peroxidase precursor, putative, expressed | Class-III peroxidase |
| Q6EUS1 | LOC_Os02g14160 | Prx27; peroxidase precursor, putative, expressed | Class-III peroxidase |
| A0A0P0VCV3 | LOC_Os01g72130 | Os01g0949750 protein; glutathione S-transferase, putative, expressed (OsGSTU5) | Tau class GST |
| Q5JKX6 | LOC_Os01g72120 | Glutathione S-transferase, putative, expressed (OsGST7) | Tau class GST |
| Q7XDA6 | LOC_Os10g34020 | Glutathione S-transferase, N-terminal domain containing protein, expressed (OsGST47) | Tau class GST |
| Q8W2T8 | LOC_Os10g22070 | Glutathione S-transferase, N-terminal domain containing protein, expressed (OsGST33) | Tau class GST |
| Q93VX8 | LOC_Os01g27480 | Glutathione S-transferase, putative, expressed (OsGST13) | Phi class GST |

**Figure S6: STRING analysis reflecting potential interacting partners of OsNUOR. (A)** Plot showing the interaction of OsNUOR (Q8H5X6) with partners in STRING database version 11.5. **(B)** The predicted interacting partners are tabulated.

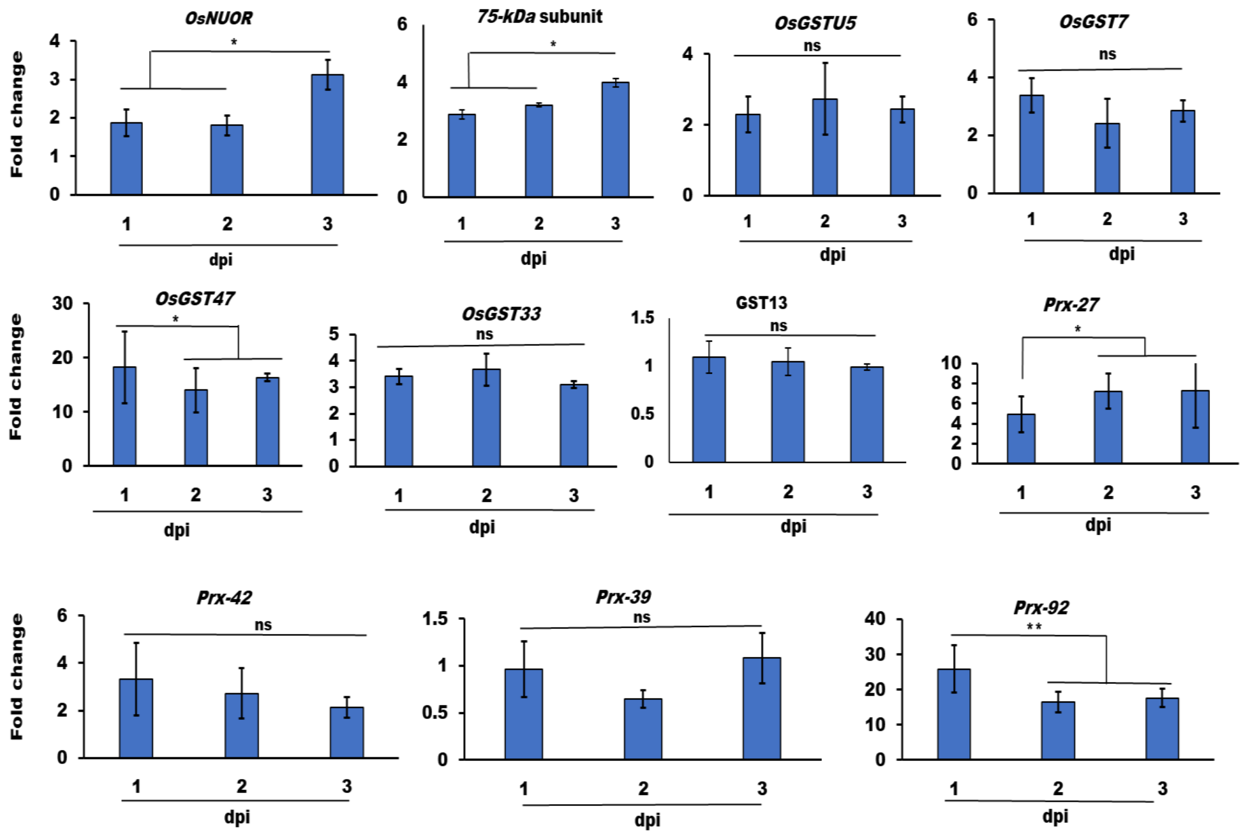


**Figure S7**: **The interacting partners of OsNUOR are upregulated under *R. solani*-infected conditions in rice.** qRT-PCR-based estimation of fold change in gene expression, with respect to 0 dpi samples, using rice *actin* gene as endogenous control. The bar denotes the average of three independent biological replicates and the error bar represents mean ± standard error. '*' indicate significant differences at $p < 0.05$ (estimated using one-way ANOVA and Student-Newman-Keuls test). ns indicates no significant difference.

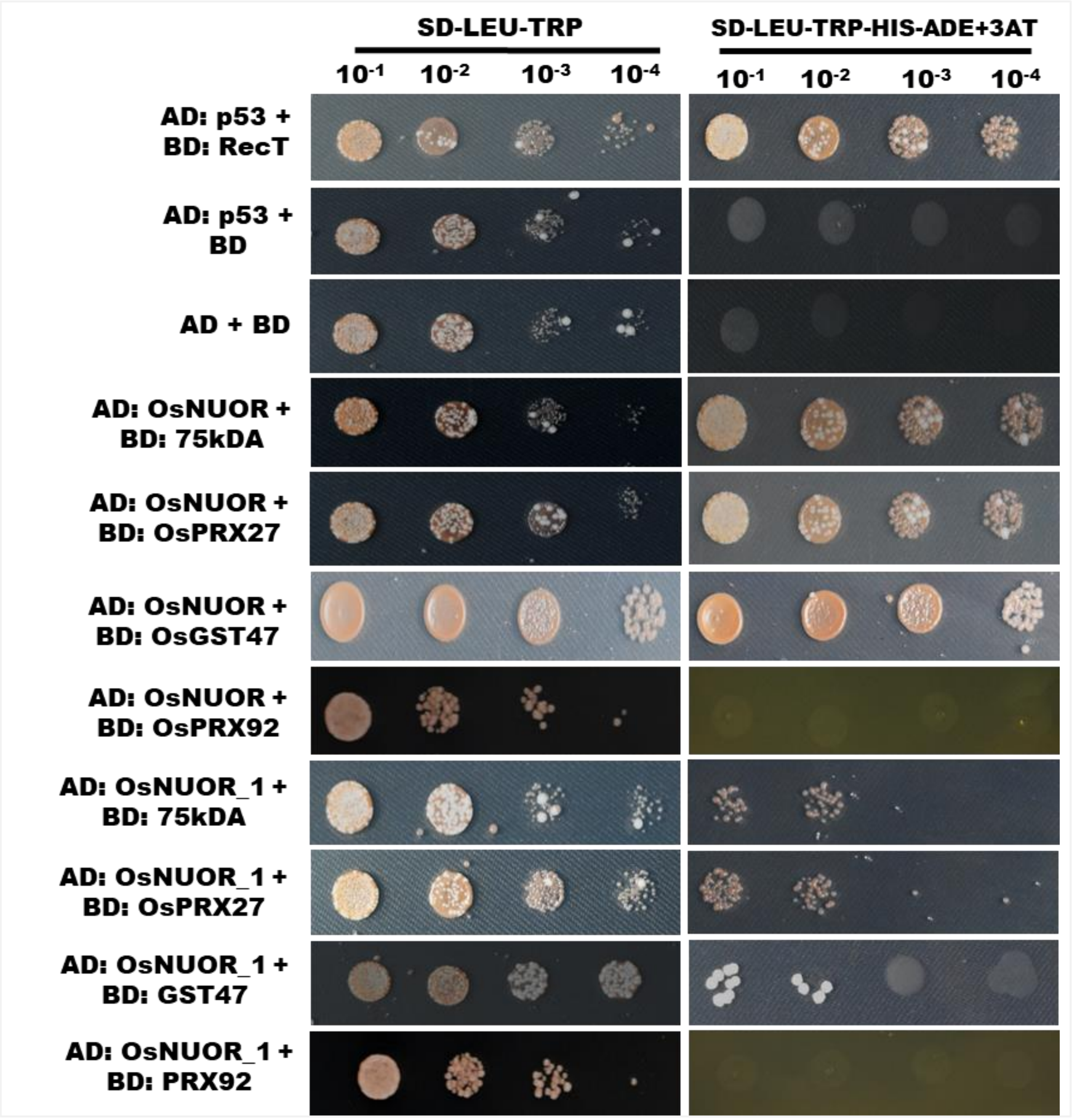


**Figure S8: Yeast two-hybrid assay to test the interaction of OsNUOR with predicted target proteins**. Y2H assay reflecting the interaction of OsNUOR/OsNUOR_1 with target proteins (75 kDa subunit of complex-1, Prx27, and GST47). The growth of yeast cells on quadruple drop-out plates (SD-Leu-Trp-Ade-His), reflects positive interactions. AD is prey vector (pGADT7) and BD is bait vector (pGBKT7). 3-AT; 3-aminotrizole (15 mM). AD: p53 + BD: RecT is used as positive control and AD: p53 +BD is used as negative control.

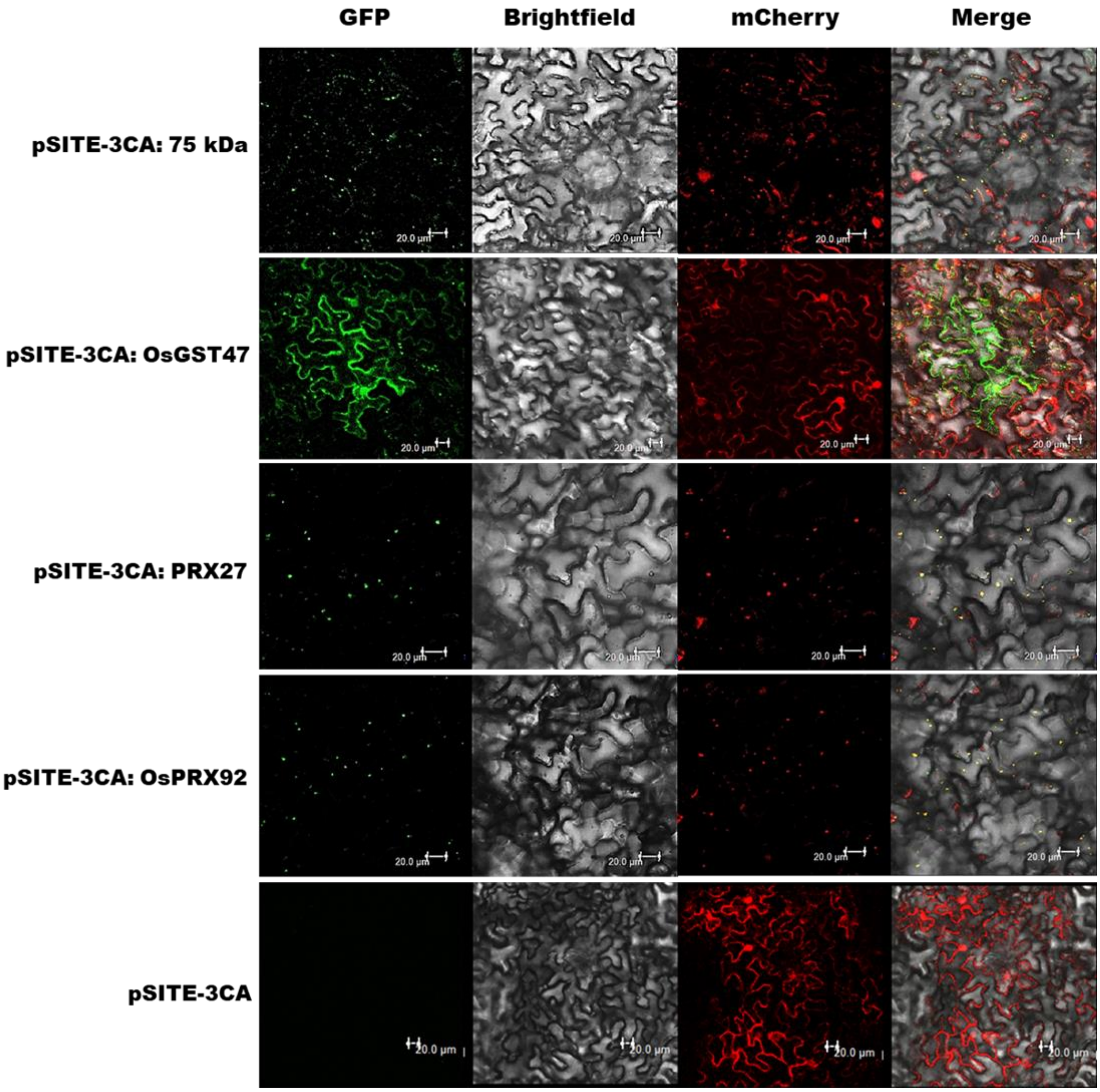


**Figure S9: Localization of selected interacting partners of OsNUOR in *N. benthamiana***. Confocal images showing localization of YFP-tagged OsGST47 in the cytosol and YFP-tagged OsPRX27, OsPRX92 and Os75 kDa subunit of complex-1 in mitochondria (being colocalized with mCherry-tagged mitochondrial marker, mt-rk CD3-991). The YFP was visualized under GFP (green fluorescent protein) filter, while mCherry was visualized under RFP (red fluorescent protein) filter. Scale bars = 20 μm.

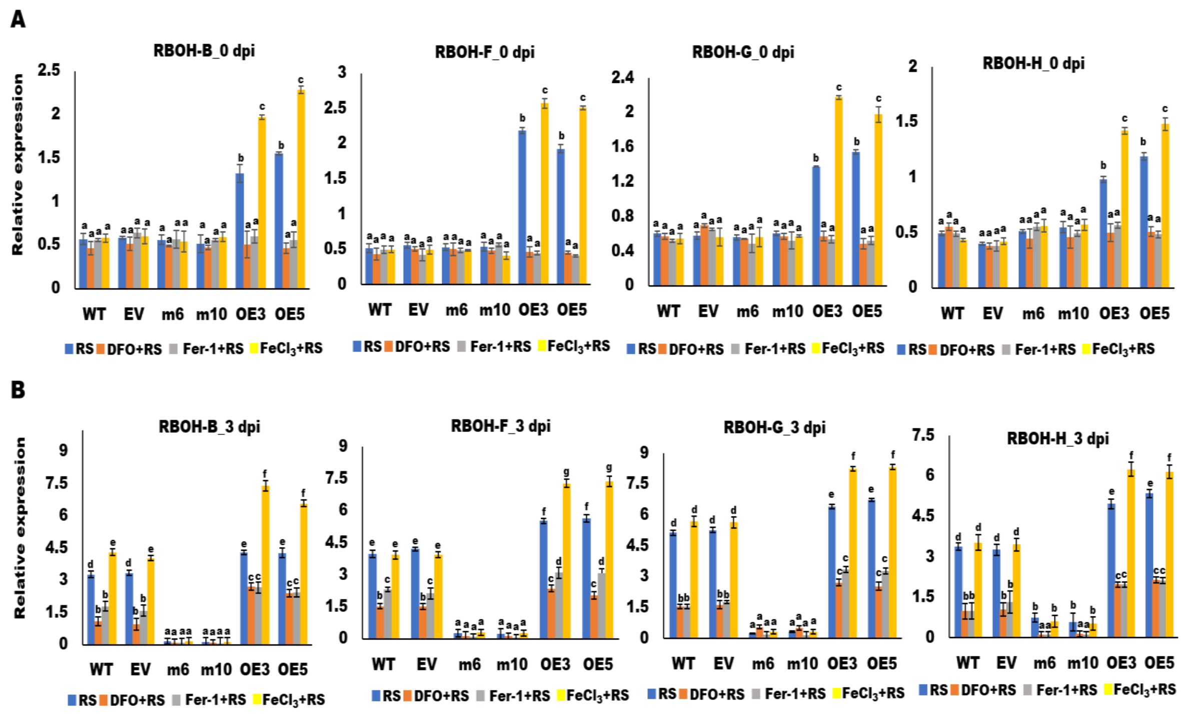


**Figure S10: Editing of *OsNUOR* compromises the expression of ROS-responsive genes.** Relative expression of *RBOH* homologs (*OsRBOH-B*, *OsRBOH-F*, *OsRBOH-G* and *OsRBOH-H*) in different rice lines with and without DFO/FER-1/$FeCl_3$ treatment at **(A)** 0 dpi and **(B)** 3 dpi of *R. solani* infection. Relative expression was quantified using *actin* gene of *Oryza sativa* as an endogenous control. The bar denotes the average of three independent biological replicates and the error bar represents mean ± standard error. Different letters represent a significant difference between samples at $p < 0.05$ (one-way ANOVA, Student-Newman-Keuls test).

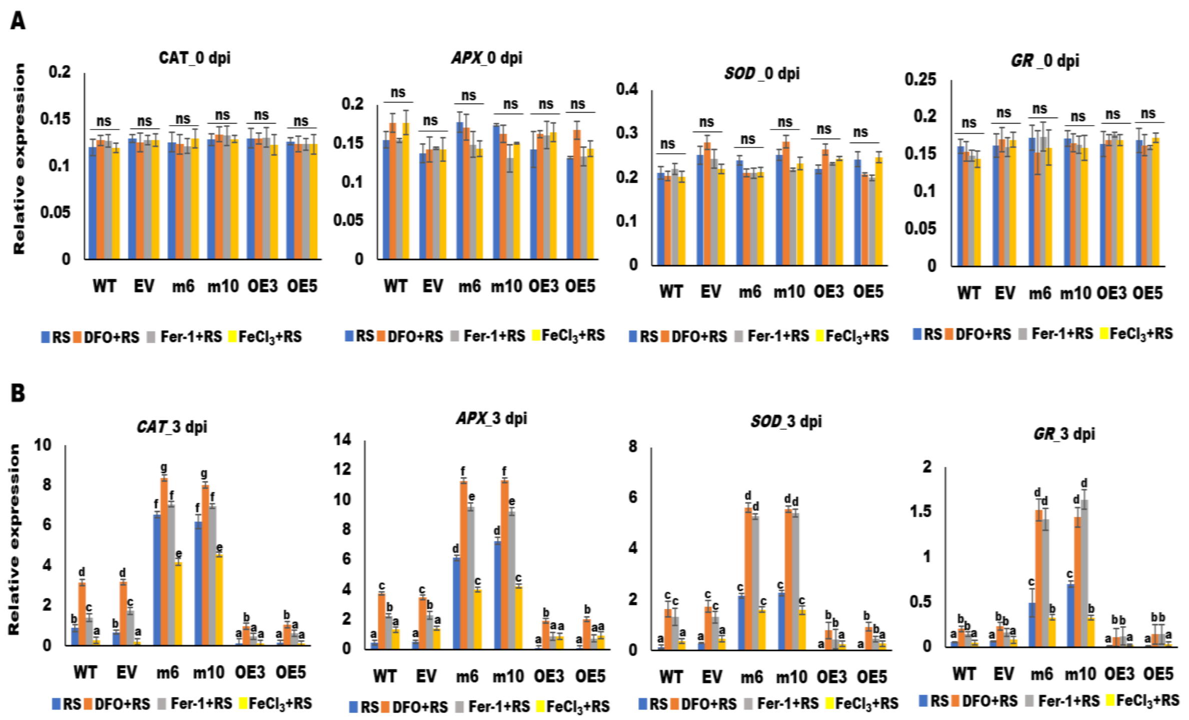


**Figure S11: Antioxidant defense is induced in edited lines, under *R. solani*-infected conditions**. Relative expression of antioxidant genes (*OsCAT*, *OsAPX*, *OsSOD* and *OsGR*) in different rice lines, with and without DFO/FER-1/$FeCl_3$ treatment at **(A)** 0 dpi and **(B)** 3 dpi of *R. solani* infection. Relative expression was quantified using rice *actin* gene as an endogenous control. The bar denotes the average of three independent biological replicates and the error bar represents mean ± standard error. Different letters represent a significant difference between samples at $p < 0.05$ (one-way ANOVA, Student-Newman-Keuls test). ns indicates no significant difference.

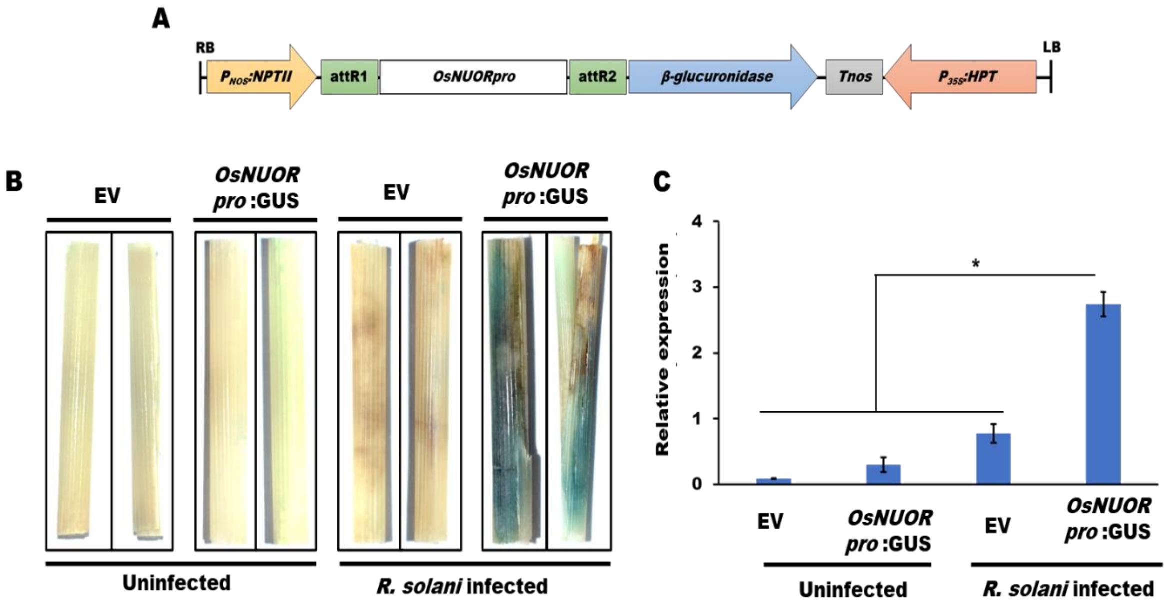


**Figure S12: *OsNUOR* is upregulated upon *R. solani* infection in rice. (A)** The map of pGWB3 gateway vector used for cloning of *OsNUOR* promoter for reporter GUS assay. **(B)** β-glucuronidase (GUS) assay reflecting activation of *OsNUOR* promoter (appearance of blue colour) in rice sheaths upon *R. solani* infection at 3 dpi. **(C)** qRT-PCR-based quantification of β-glucuronidase (*GUS*) gene expression using rice *actin* gene as an endogenous control. The bar denotes the average of three independent biological replicates and the error bar represents mean ± standard error. '*' indicate significant differences at $p < 0.05$ (estimated using one-way ANOVA and Student-Newman-Keuls test).

**Supplementary table 1: TargetP and SignalP results of *OsNUOR* and its interacting partners**

| Protein | Uniprot ID | TargetP | SignalP | Likelihood |
|---|---|---|---|---|
| OsNUOR | Q8H5X6 | Mitochondrial transfer peptide | - | 0.6528 |
| Mitochondrial complex-I 75-kDa subunit | Q8W317 | Mitochondrial transfer peptide | - | 0.4859 |
| Prx27 | Q6EUS1 | - | Secretory | 0.9804 |
| OsGST47 | Q7XDA6 | Other | - | 0.7265 |
| PRX92 | Q5Z8H7 | - | Secretory | 0.9944 |

**Supplementary table 2: List of primers used in the study**

| Gene Id | Primer name | Primer sequence (5'→ 3') (Highlighted region represents overhang sequence for directional cloning) |
|---|---|---|
| **For full length gene cloning (localization, BiFC, Y2H, deletion construct)** | | |
| LOC_Os07g37730 | OsNUOR-FP | CACCATGGCGTGGTCGAGGATT |
| | OsNUOR-RP | GCCGATCCTGGTGTTGTC |
| | OsNUOR_1-FP | CACCATGGCGGCGGCGGTGCCGGAC |
| | OsNUOR_1-RP | GCCGATCCTGGTGTTGTCCCTGCCGAACA |
| LOC_Os03g50540 | 75 kDa-FP | CACCATGGCCTTCTTCGCGAGGGCGATC |
| | 75 kDa-RP | CTTCTTCAACAAGGTTGCACTGCA |
| LOC_Os02g14160 | OsPRX27-FP | CACCATGGCTGGAGCCGTGCTGGTGA |
| | OsPRX27-RP | GTTGGGCTTCCGGCAGTTGGAGC |
| LOC_Os10g34020 | OsGST47-FP | CACCATGCGGGTGATGGTGGCGCTGA |
| | OsGST47-RP | AATCAGCTTAGCTGTGAACTCGACC |
| LOC_Os06g48010 | OsPRX92-FP | CACCATGAGGAGCTTCCATTTCGTCGTCGT |
| | OsPRX92-RP | GTTGGGGAACCTGCAGTCGCG |
| **For guide RNA cloning (Highlighted region represents flanking sequence)** | | |
| LOC_Os07g37730 | gRNA1-FP | TAGGTCTCCGAGGATTGCGAGGTTTTAGAGCTAGAA |
| | gRNA1-RP | CGGGTCTCACCTCGACCACGCTGCACCAGCCGGG |
| | gRNA2-FP | TAGGTCTCCCGAGCTGAGCGAGTTTTAGAGCTAGAA |
| | gRNA2-RP | CGGGTCTCACTCGATCTCCCTTGCACCAGCCGGG |
| **For PCR confirmation** | | |
| OQ579018.1 | Cas9-FP | AGTTCATCAAGCCCATCCTG |
| | Cas9-RP | GAAGTTTCTGTTGGCGAAGC |
| AB303069.1 | HPT-FP | GATGTAGGAGGGCGTGGATA |
| | HPT-RP | ATTTGTGTACGCCCGACAGT |
| LOC_Os01g16414 | OsActin1-FP | ACAGTTCAACCCTCCAGCAC |
| | OsActin1-RP | TGATTGTGCTGTCCCAGAAG |
| X04879.1 | CaMV35S-FP | ACTATCCTTCGCAAGACCCTT |
| LOC_Os07g37730 | OsNUOR-RP | GCCGATCCTGGTGTTGTC |
| **For amplification of target region for sequencing** | | |
| LOC_Os07g37730 | OsNUOR-UTR-FP | CTTTGAATTGATCTGCAGGTTCTC |
| | OsNUOR-g1-RP | AGGAGGCGAGGAAGAAGTAGG |

| | | |
|---|---|---|
| | OsNUOR-g2-FP | GCGTGGTGAAGGAGGTGAA |
| | OsNUOR-g2-RP | TCAGCCGATCCTGGTGTTG |
| LOC_Os01g61410 | 61410NUOR-g1-FP | TCCAGATCTCAGCCTCTCGG |
| | 61410NUOR-g1-RP | CAGCAAAGCCCAGAAATCCG |
| | 61410NUOR-g2-FP | GCCTGGAAAAGAGGCATGGTA |
| | 61410NUOR-g2-RP | GACACCTGTTGGACCACCTC |
| **For qRT-PCR analysis** | | |
| XM_015784227.3 | OsActin-FP | CCGAGACCTCACTGACCATC |
| | OsActin-RP | TAATCAAGGGCGACATAAGC |
| MK481078.1 | Rs18S-FP | CCCCCTGTGCACTTGTGAG |
| | Rs18S-RP | ATCGCATTTCGCTGCGTTC |
| LOC_Os07g37730 | OsNUOR-FP | AGCTCAGCGACTTCATCACC |
| | OsNUOR-RP | TACTTGGAGAGGTGGTCCGT |
| LOC_Os01g61410 | 61410NUOR-FP | CGTGGTGTTACTGACCATGC |
| | 61410NUOR-RP | TGCGCTTTTCTTCCTCCGAT |
| LOC_Os03g50540 | Os75kDa-FP | ACAGTGATGACCCGTTGCAT |
| | Os75kDa-RP | CCACTGCCACGACCTAACAT |
| LOC_Os01g72130 | OsGSTU5-FP | GGAGATTCTTCGGAGGCGAC |
| | OsGSTU5-RP | CCAAGGTCACCCCACATACC |
| LOC_Os01g72120 | OsGST7-FP | ACAACCCCGTCCACAAGATG |
| | OsGST7-RP | CGACGTACTCGACGATGACC |
| LOC_Os03g25280 | OsPRX42-FP | AGATCAACGTCCCAGGTGGA |
| | OsPRX42-RP | GTTCACGTTGAAGTTGGGCG |
| LOC_Os03g13200 | OsPRX39-FP | AGATCTCCGTCGAACTGTCC |
| | OsPRX39-RP | GGGCTGGGGAGGTTATTGTC |
| LOC_Os06g48010 | OsPRX92-FP | AGGGGTTCACGACGGTGATA |
| | OsPRX92-RP | TCCACTCAGGAAGACGGAGT |
| LOC_Os02g14160 | OsPRX27-FP | TTCCACGACTGCTTCGTCAA |
| | OsPRX27-RP | GTTCTTCTCGCCGGTGATCG |
| LOC_Os10g34020 | OsGST47-FP | CGCTTCTGGGGACAGTACAT |
| | OsGST47-RP | GAGCGGTTGTCATTTCGTCG |
| LOC_Os10g22070 | OsGST33-FP | GAACCCTAGCCCGTTCGTC |
| | OsGST33-RP | CGAGGAGGAGGTCGGTCTT |
| LOC_Os01g27480 | OsGST13-FP | AAACCCGTTTGGGCAAGTTC |
| | OsGST13-RP | CTTCCGCAGTATGTACCGGC |
| LOC_Os01g52500 | OsNADP-ME2-FP | TGGTTCATTGTCGGAGCACA |
| | OsNADP-ME2-RP | TTCTTGCGGCACTCCTCAAT |
| LOC_Os01g25820 | OsRBOHB-FP | CTCGCTGTTCCTGTTGTCCT |
| | OsRBOHB-RP | CCGGGATACACTGCAACCTT |
| LOC_Os05g38980 | OsRBOHD-FP | GTTCAGGAGGAGCGTTGTGA |

| | | |
|---|---|---|
| | OsRBOHD-RP | CAGTAGGTGATGGGCGTACC |
| LOC_Os08g35210 | OsRBOHF-FP | GATCGTCATGGTGGTGCTCA |
| | OsRBOHF-RP | CGAGTACCAGAAGGCGTTGA |
| LOC_Os09g26660 | OsRBOHG-FP | GGGTTCAACGCCTTCTGGTA |
| | OsRBOHG-RP | AAATACCAGCGACGGACGAG |
| LOC_Os12g35610 | OsRBOHH-FP | TACTCCCACCACCTCTTCGT |
| | OsRBOHH-RP | CTTCCACCAAGTCCGGTTGA |
| AB289766.1 | Gus-FP | AGACTGTAACCACGCGTCTG |
| | Gus-RP | TTGTCCAGTTGCAACCACCT |
| Os06t0185900-01 | OsGPX4-FP | TGCACAAGAACCTGGGTCAA |
| | OsGPX4-RP | TGTGTTAGGCCCGTTGACAT |
| Os11t0284900-01 | OsGPX5-FP | CCGCACCGCTATACAAGTTC |
| | OsGPX5-RP | CGCTCAACAACCTTGCCATC |
| NM_001401748.1 | OsCAT-FP | GCCAGGTACGTCAAGTTCCA |
| | OsCAT-RP | GAGTCGTAGAGGTCCTGGGT |
| NM_001402136.1 | OsAPX-FP | CCCAGGAAGGGAGGACAAAC |
| | OsAPX-RP | CGAAGACCTGCCTTAGGTGG |
| NM_001402510.1 | OsMnSOD-FP | GTCGCCAACTACAACAAGGC |
| | OsMnSOD-RP | CGTTGAACTTGATGGCGCTC |
| XM_015772375.2 | OsGR-FP | TTTCTCCCAACCACCCATCG |
| | OsGR-RP | TTGCCCTAAGTGGCCTGAAG |
